# Airflow Constraints Limit Natural Airborne Transmission of Tuberculosis

**DOI:** 10.1101/2025.10.27.684829

**Authors:** Kubra F. Naqvi, Yuhui Guo, Yash Kulkarni, Deepak Sapkota, Pei Lu, Shibo Wang, Beatriz R. S. Dias, Victoria A. Ektnitphong, Arabella E. Martin, Bret M. Evers, Lenette L. Lu, Hui Ouyang, Lydia Bourouiba, Michael U. Shiloh

## Abstract

Tuberculosis (TB) is transmitted through the air, yet the determinants of natural airborne transmission remain poorly defined. Early twentieth-century guinea pig studies demonstrated efficient airborne transmission of *Mycobacterium tuberculosis* (Mtb), but this paradigm has not been reestablished in contemporary containment facilities. Here, we show that ventilation can impose airflow constraints that suppress transmission under otherwise permissive conditions. Using a guinea pig model of animal-to-animal exposure, we combined transmission experiments with quantitative airflow measurements and particle transport modeling to explain why some housing configurations fail to support effective exposure. Static environments and excessive unidirectional airflow prevented transmission, whereas controlled low-velocity airflow restored evidence of exposure, including tuberculin skin test conversion, antigen-specific immune responses, and pulmonary inflammation consistent with early infection. These findings identify airflow as a critical constraint on airborne TB transmission and provide a reproducible framework to dissect host, microbial, and environmental determinants of spread.

## INTRODUCTION

Tuberculosis (TB), caused by infection with *Mycobacterium tuberculosis* (Mtb), remains one of the leading causes of infectious mortality worldwide (*1*). Interrupting the transmission cascade is essential to reducing global incidence, yet progress has been hindered by an incomplete understanding of the host, bacterial and exposure-related factors that govern airborne transmission. Experimental systems that faithfully recapitulate natural transmission are therefore critical for dissecting the biological determinants of infection and for accelerating the development of transmission-blocking interventions and vaccines.

Guinea pigs have played a central role in the history of TB research (*2*). In 1882, Robert Koch established their use as a model for TB pathogenesis, noting that naïve animals housed near infected ones occasionally developed spontaneous pulmonary disease (*3, 4*). In the 1920s and 1930s, David Perla and Max Lurie expanded on these observations, demonstrating that healthy guinea pigs co-housed with infected animals developed bronchogenic TB, and that transmission could occur even between animals housed in separate cages within the same room (*5–9*). These seminal studies provided the first direct experimental evidence for airborne transmission of Mtb. Decades later, Riley, Wells, and Mills extended this concept to humans by showing that air exhausted from TB wards could infect sentinel guinea pigs (*10–14*).

Despite the foundational impact of these studies, the experimental paradigm of natural airborne transmission was not widely carried forward into the modern biosafety era. Contemporary biosafety level 3 (BSL-3) facilities impose stringent containment and ventilation requirements that differ substantially from the environments in which early transmission experiments were conducted. How these modern conditions influence the biological feasibility of natural airborne transmission has received relatively limited experimental attention.

Recent studies using alternative animal models, including minipigs (*15*) and ferrets (*16*), have demonstrated animal-to-animal transmission of Mtb, but these systems require high infectious doses and do not fully capture natural respiratory behaviors such as coughing. The guinea pig remains well suited for studying transmission biology: it is relatively low-cost compared to other large-animal models, develops necrotizing granulomas that resemble human disease (*17*), harbors Mtb within its airways during infection (*18*), and exhibits respiratory neurophysiology closely resembling that of humans (*19, 20*). While guinea pigs have long been used as sentinels for human-to-animal transmission in air sampling studies (*12, 13, 21–23*), a modern system that supports natural guinea pig-to-guinea pig airborne transmission under BSL-3 conditions has not been established. This gap raises the possibility that features of contemporary cage-level airflow and ventilation may impose constraints on transmission that are independent of host susceptibility or bacterial virulence.

Airborne transmission of Mtb depends on the generation, persistence, and inhalation of pathogen-containing aerosols produced during normal respiratory behaviors, including coughing, breathing, sneezing, and singing (*24–35*). Once expelled, these aerosols are shaped by environmental factors such as temperature, humidity, and ventilation (*33*) that influence particle desiccation, dispersal, and persistence (*36–38*). In confined animal housing systems, effective biological exposure is therefore determined not only by aerosol generation but also by airflow patterns within the exposure environment. Ventilation at the level of the cage can influence whether aerosols are retained long enough to reach a susceptible host or are removed before inhalation. At the same time, under modern BSL-3 containment, negative-pressure ventilation imposed at the facility level can interact with imperfectly sealed enclosures, potentially altering airflow within cages in ways that are not apparent from cage design or nominal ventilation rates alone. Together, these considerations raise the possibility that multiple, interacting airflow constraints shape biological exposure during experimental transmission, yet their combined impact has been difficult to define in living systems.

To establish a modern experimental system capable of supporting natural airborne transmission, we revisited guinea pig-to-guinea pig spread of Mtb using housing designs adapted for contemporary BSL-3 environments. Our initial goal was to establish a functional animal-to-animal transmission model that preserved biological exposure while meeting biosafety requirements, and to understand which factors proved limiting as the system was developed. When early housing configurations failed to support transmission, we iteratively modified cage design and airflow conditions and incorporated quantitative particle measurements to assess how features of the exposure environment influenced aerosol retention and inhalation. Computational modeling was then used to interpret these observations and explain how airflow patterns and enclosure design shape access to the recipient breathing zone. Together, this stepwise biological and engineering approach revives a classical transmission paradigm with modern tools and establishes a biologically grounded platform for dissecting host, microbial, and exposure-related determinants of airborne tuberculosis transmission.

## RESULTS

### Initial housing configurations do not support guinea pig-to-guinea pig airborne Mtb transmission under BSL-3 conditions

Guided by early transmission studies in guinea pigs (*8*), we first tested whether a simple co-housing design could support natural airborne transmission under modern BSL-3 conditions. We therefore tested the first iteration of our guinea pig transmission system (system 0) by co-housing naïve recipient guinea pigs with Mtb-infected donors, separated by a perforated plexiglass barrier for 12 months (Fig. 1A, B). Infection of naïve animals was monitored every other month by tuberculin skin testing (TST).

**Figure 1:**
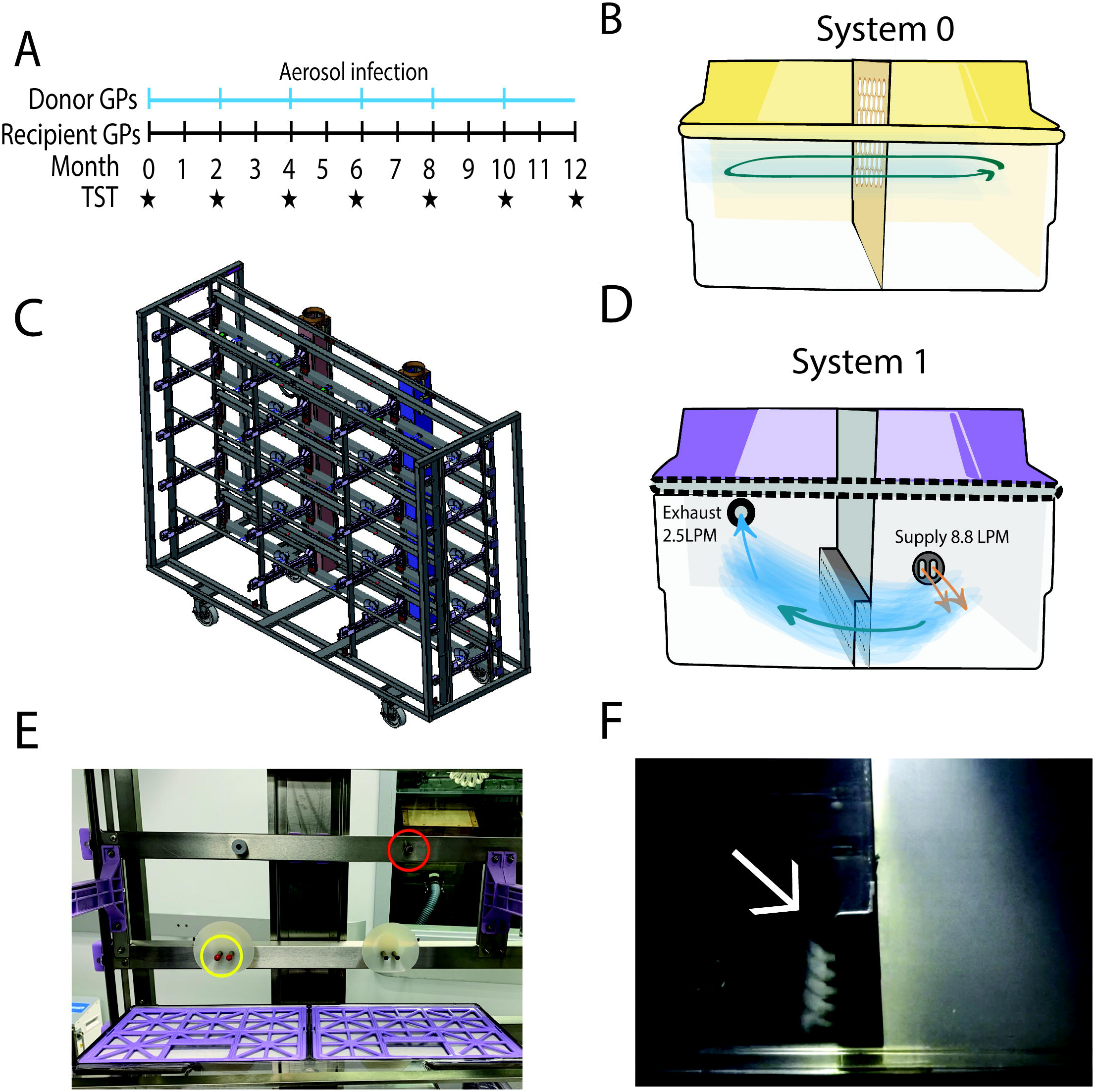
Airflow Modifications of Preliminary Transmission Systems. (A) Experimental scheme for 12-month transmission studies. Blue ticks indicate when new donor groups were added to cages and stars denote when TSTs were performed. (B) System 0 cage set up with static airflow and plexiglass divider between donor and recipient compartments. (C) 15 cage rack as available by Allentown, Inc. (D) System 1 cage set up. Donor side is connected to supply hose which delivers 8.8 L/min into the cage. Air then passes through a stainless-steel divider to recipient compartment and is exhausted at 2.5 L/min. (E) Modifications to cage rack include plugging the exhaust port on the donor side and supply port on the recipient side to drive unidirectional airflow. (F) Smoke test to confirm unidirectional movement of air within system 1. See also Supplemental Movie 1.

This initial design emphasized simplicity, low-cost and ease of access to system components. We modified an existing static cage by inserting a plexiglass divider containing large holes to separate donor and recipient animals (Fig. 1B). The cages were housed on metal shelving located in the most negatively pressurized room of our ABSL3 facility (relative negative pressure −0.06 WC). Because guinea pigs are highly susceptible to progressive tuberculosis following low-dose infection (50-100 CFU) (*17*), infected donors were replaced every 7-8 weeks to maintain a constant infectious source. After 12 months of exposure, none of the recipient animals converted their TSTs or showed clinical or histopathologic evidence of infection, and no Mtb CFU were recovered at necropsy. The absence of transmission in this initial configuration indicated that this simple co-housing design was insufficient to support guinea pig-to-guinea pig airborne transmission under modern BSL-3 conditions.

The absence of transmission under these conditions contrasted sharply with historical reports from Perla and Lurie and suggested that modern biosafety engineering constraints might fundamentally alter airflow compared to the open systems used a century ago. In contemporary BSL-3 facilities, negative-pressure ventilation ensures containment but may also disrupt airborne particle movement between animals. On this basis, we next tested whether enforcing directional airflow from infected donors to naïve recipients would be sufficient to permit transmission.

To test this, we modified a commercially available guinea pig housing system (Allentown, Inc.) where cages are connected to both supply and exhaust air blowers to create “system 1” (Fig. 1C, D). The new design incorporated a stainless-steel barrier between donor and recipient animals perforated with two panels of holes positioned at the typical nose and mouth height of an adult guinea pig. To establish unidirectional airflow, we blocked the donor-side exhaust port (right) and the recipient-side supply ports (left) with rubber caps (Fig. 1E). Smoke testing confirmed right-to-left airflow from the donor to the recipient compartment (Fig. 1F, Supplemental Movie 1). Recipient animals were then exposed to infected donors for 12 months under these unidirectional airflow conditions. Despite these adjustments, no transmission occurred: all recipients remained TST-negative and free of histopathologic or microbiologic evidence of infection. These findings indicated that unidirectional airflow alone was insufficient to overcome transmission constraints imposed by BSL-3 conditions.

### Airflow and cage design determine aerosol particle retention

Because systems 0 and 1 did not support transmission, we next directly measured aerosol particle transport across housing configurations to determine whether physical features of the cages limited retention and exposure of airborne particles. We undertook an iterative series of aerosol studies to define the parameters that would permit particle transport from donor to recipient compartments.

To test aerosolized particle transport, we connected prototype cages to a supply air line on the donor side and an exhaust vacuum on the recipient side, allowing controlled regulation of airflow within the system. To simulate infectious particles, we used a particle generator to produce aerosols ranging from 0.8-7 µm and sampled air at three locations: upstream (before entering the cage), within the donor compartment, and within the recipient compartment (Fig. 2A). Particles were loaded for 1 hour to establish steady-state size distribution (Supplemental Fig. 1A) and sampled three times for 20 seconds totaling 1 minute at each position.

**Figure 2:**
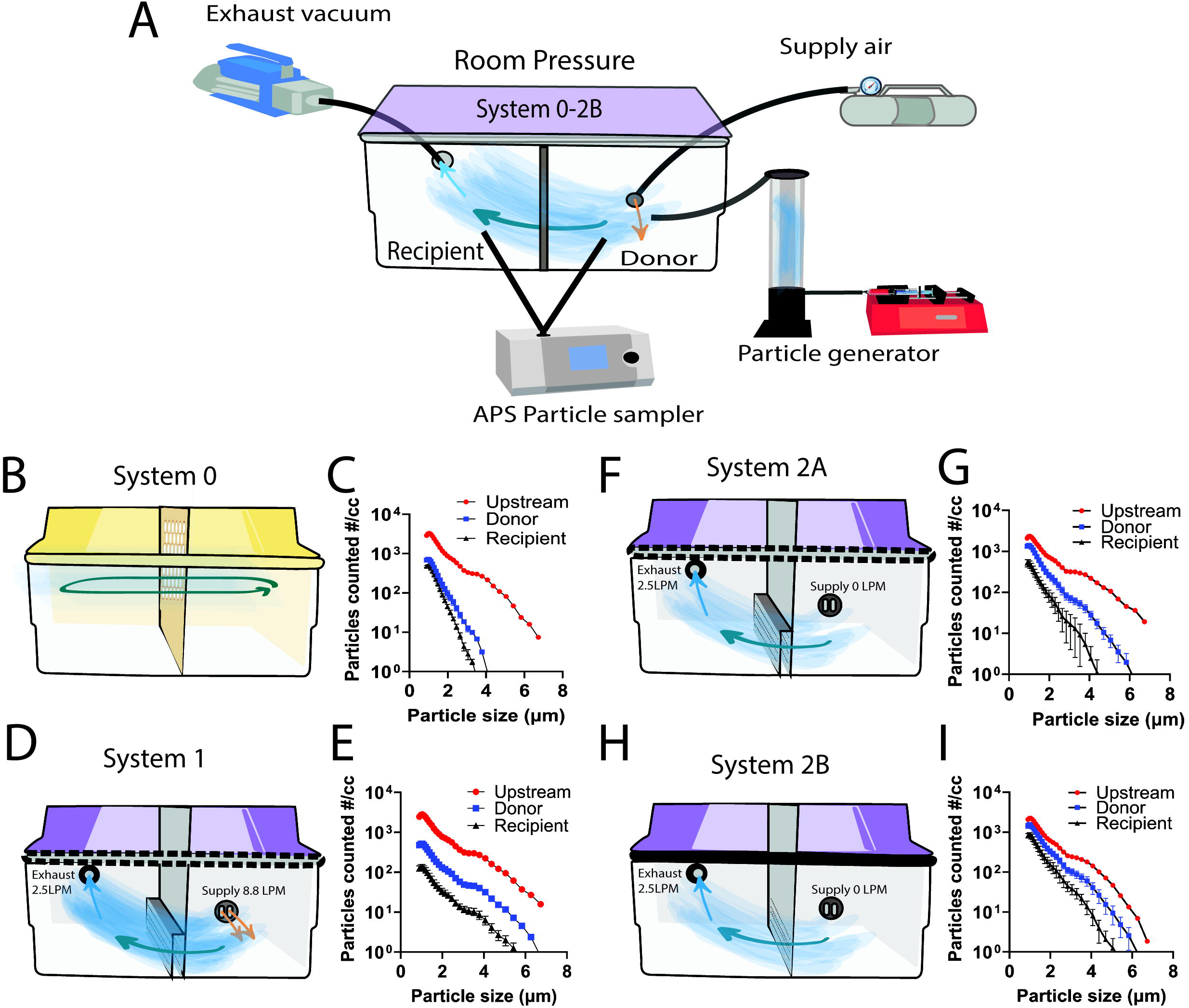
Particle movement driven by airflow under ambient pressure. (A) Experimental overview of particle transport. Cage systems are connected to supply air (where applicable) and exhaust vacuum to control airflow within the cages. Particles are introduced to the donor compartment using a particle generator. An APS particle sampler collects particles upstream before entering the cage, in the donor and recipient compartments. (B-C) Schematic and particle sampling of System 0 under static flow conditions. (D-E) Schematic and particle sampling of System 1 with supply air set to 8.8 L/min and exhaust to 2.5 L/min. Dashed line indicates unsealed cage lid. (F) System 2A schematic with air supply removed (0 L/min) and exhaust set to 2.5 L/min. (G) Particle transport within system 2A. (H-I) Schematic and particle transport of system 2B modified to include cage lid sealing (solid line) and a single stainless-steel barrier. All particle transport results are represented as number of particles per cubic centimeter of air (#/cc) in the size range of 0.8-7 µm. Sampling of particles includes upstream (red) before entering the cage, in the donor compartment (blue) and the recipient compartment (black).

We first characterized particle behavior under ambient room pressure, beginning with the earliest systems, 0 and 1. In system 0, which lacked directional airflow, we observed marked particle loss from both donor and recipient compartments, particularly for particles larger than 4 µm (Fig. 2B-C). System 1, operated with the manufacturer’s default airflow settings (8.8 L/min supply; 2.5 L/min exhaust), also showed significant particle depletion (Fig. 2D-E). These findings indicated that the high ventilation rates used for standard animal husbandry, intended to prevent excessive humidity, ammonia and CO_₂_ accumulation, were incompatible with retention of aerosolized particles. Successful transmission would likely require precise titration of airflow to maintain particle dwell time without compromising animal welfare.

Guided by these results, we next developed system 2A, in which air movement was driven solely by the exhaust fan (2.5 L/min). This modification improved particle retention within both donor and recipient compartments under ambient conditions, especially for particles less than 2 µm (Fig. 2F-G).

Inspection of system 1 also revealed a structural limitation: its double-walled stainless-steel barrier created a wide and obstructed path for particle transfer between compartments. To address this, we engineered a new single-walled stainless-steel barrier with larger perforations to reduce resistance to airflow. We also sealed the cage to minimize particle leakage by taping the lid with aluminum tape, adding a rubber gasket along the lid-base junction, and securing the lid with four spring-loaded clamps. The resulting configuration, designated system 2B, was designed to optimize both airflow directionality and particle retention (Fig. 2H). Under ambient pressure, system 2B achieved particle retention comparable to system 2A, with stable concentrations of particles <2 µm in both compartments (Fig. 2I). The similarity between the two systems suggested that the improvements achieved by airflow titration and cage sealing were robust across minor design variations. However, because all prior experiments had occurred under negative-pressure BSL-3 conditions, further testing under those laboratory containment conditions was essential to evaluate real-world performance. Together, these measurements demonstrated that standard housing configurations and ventilation rates used for animal care rapidly deplete aerosolized particles, providing a physical explanation for the absence of transmission observed in earlier system iterations.

### Negative-pressure BSL-3 conditions alter aerosol particle retention across housing configurations

Because all Mtb transmission studies must be performed under BSL-3 conditions, we next evaluated how negative pressure, an essential safety feature, affects particle retention within the cage transmission systems. To do this, we placed each system inside a custom airtight chamber maintained at the negative-pressure setting used in our ABSL-3 facility (relative negative pressure −0.03 WC) (Fig. 3A). Of note, the negative pressure imposed on the system mimicked the settings of the room in which the transmission studies were performed, −0.03 (system 1, 2A, 2B) or −0.06 (system 0). Under these conditions, we observed a striking loss of aerosolized particles in both donor and recipient compartments of system 0 (Fig. 3B, C). The unsealed cage design allowed particles to escape through lid gaps and barrier edges, resulting in near-complete depletion of particles across all measured size ranges. System 1 also exhibited substantial particle loss under negative pressure, particularly in the recipient compartment (Fig. 3D, E), indicating that unidirectional airflow alone could not overcome the strong external flow imposed by facility ventilation. System 2A, which incorporated reduced exhaust flow and a modified barrier, also showed lower particle retention in the recipient compartment (Fig 3F, G). The most dramatic improvement was achieved with system 2B, which combined controlled exhaust flow with cage sealing. Under negative pressure-induced flow, system 2B maintained high concentrations of aerosols within both donor and recipient compartments, demonstrating markedly improved particle retention compared to all previous iterations (Fig. 3H, I).

**Figure 3:**
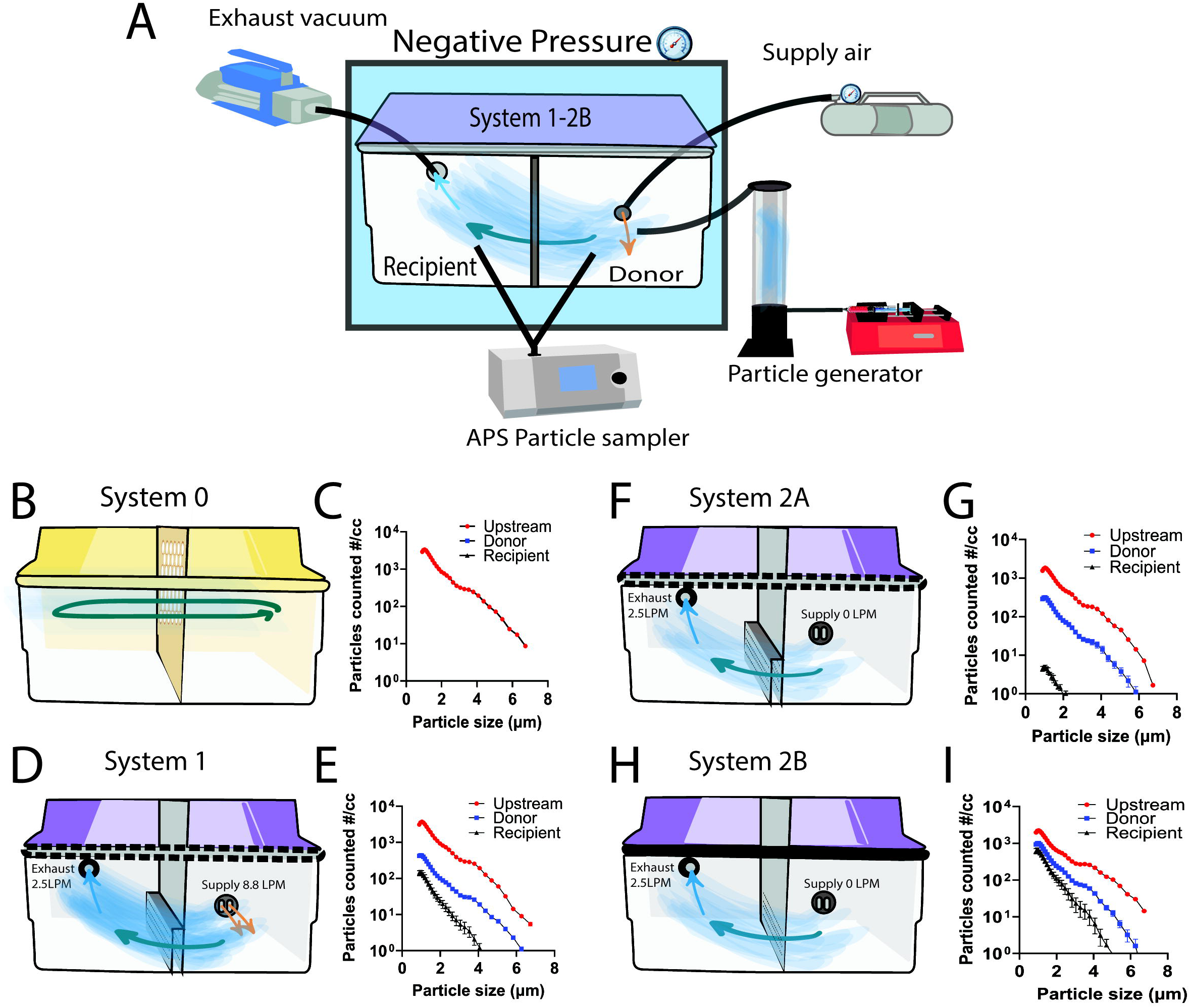
Negative pressure influences retention of particles within cage systems. (A) Experimental set-up of particle transport where cage systems are placed inside a custom chamber under negative pressure. (B-C) System 0 cage design and particle movement within cage system while under negative pressure (−0.06 WC). (D-E) Addition of unidirectional airflow in system 1 design and particle transport within system 1 under negative pressure (−0.03 WC). Dashed line indicates unsealed cage lid. (F-G) Schematic of system 2A with airflow driven by the exhaust port (2.5 L/min) and particle transport under negative pressure (−0.03 WC) conditions. (H-I) System 2B schematic with modified cage lid sealing (solid line) and single cage barrier and particle transport within system 2B. All particle transport results are represented as number of particles per cubic centimeter of air (#/cc) in the size range of 0.8-7 µm. Sampling includes upstream (red) before entering the cage, in the donor compartment (blue) and the recipient compartment (black).

Together, these results reveal that negative pressure-induced flow profoundly alters internal cage mixing, often removing aerosols before they can traverse from donor to recipient compartments. Through progressive modification of airflow, barrier design, and cage sealing, system 2B successfully counteracted these losses, providing stable particle retention even under the stringent airflow conditions required for BSL-3 containment. This optimized configuration was therefore selected for subsequent transmission experiments.

### Biological constraints define airflow regimes compatible with transmission

Because particle retention alone does not establish biological exposure, we next asked whether airflow conditions that preserved aerosols also supported bacterial transport and inhalation in vivo. To model movement of bacteria-containing particles, we nebulized a suspension of *Mycobacterium smegmatis,* a non-pathogenic mycobacterial species, in the donor compartment and measured bacterial recovery from air sampled on the recipient side. Recovery of *M. smegmatis* showed a clear dependence on airflow rate, with the highest recovery observed at the lowest exhaust setting of 0.15 L/min (Fig. 4A). This pattern suggested that reduced airflow prolongs particle dwell time within the cage and enhances the likelihood of bacterial transport between compartments.

**Figure 4.**
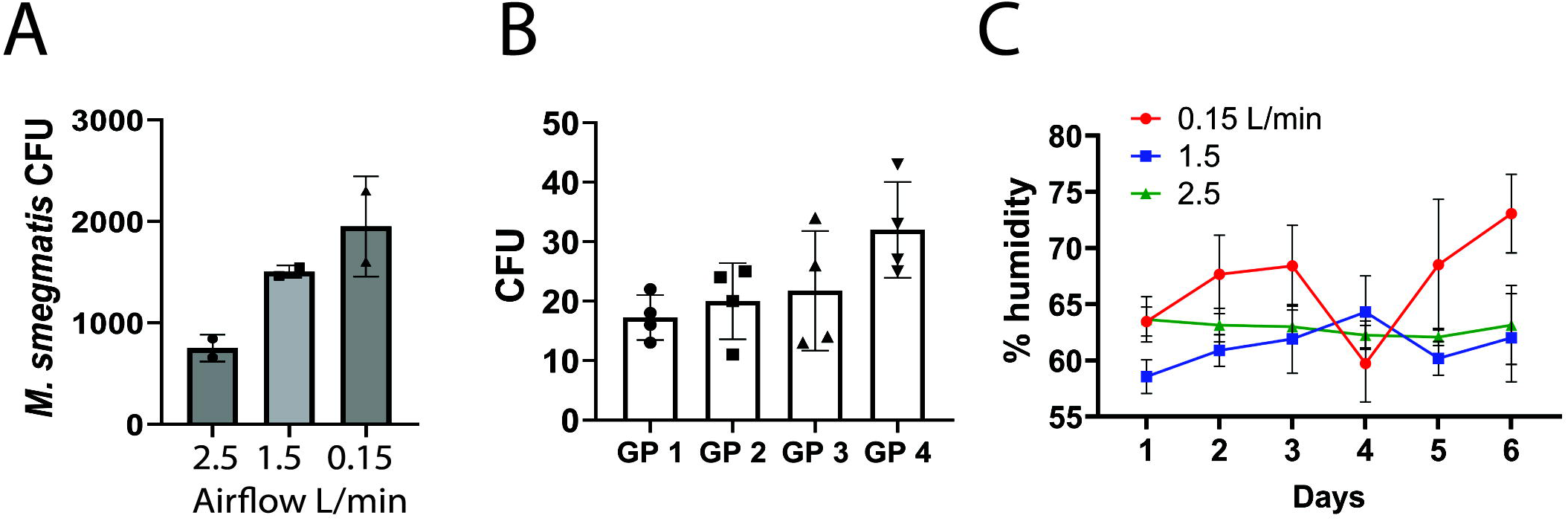
Optimization of environmental factors which influence transmission efficiency. (A) Recovery of particles containing *M. smegmatis* from the recipient compartment using a low-flow button sampler at varying exhaust flow rates (0.15, 1.5 and 2.5 L/min). (B) Nebulization of *M. smegmatis* in the donor compartment and recovery of bacteria from lungs of recipient guinea pigs in system 2B cages. Colony forming units (CFU) quantified from lung homogenates plated across four agar plates from four different guinea pigs. Each point represents CFU on an individual plate. (C) Humidity accumulation during guinea pig housing in system 2B cages. Relative humidity (%) in three system 2B cages with guinea pigs housed in each compartment (two per cage) across 6 days. Three different exhaust flowrates tested include 0.15, 1.5, and 2.5 L/min. Cages were changed at day 4 for all flowrates.

We then tested if aerosolized bacteria from the donor compartment could deposit into the lungs of naïve guinea pigs housed on the recipient side. We placed individual guinea pigs in the recipient compartment and nebulized a suspension of *M. smegmatis* in the donor compartment for one minute to simulate a cough paroxysm. After 15 minutes of exposure to *M. smegmatis*, we collected and homogenized the lungs of recipient guinea pigs and quantified colony forming units (CFU). All four recipient guinea pigs had recoverable *M. smegmatis* CFU in their lungs (Fig. 4B), demonstrating that aerosolized bacteria generated in the donor compartment were effectively transported and inhaled by recipient animals.

Because humidity strongly influences aerosol dynamics by affecting particle desiccation and settling (*33, 39–42*), we next quantified relative humidity (RH) in occupied cages. We housed two guinea pigs (one per compartment) for six days in system 2B cages, and measured RH daily at three exhaust flow rates (2.5, 1.5, and 0.15 L/min). At the lowest airflow rate, humidity rose rapidly, reaching approximately 75 percent by day six (Fig. 4C). Higher airflow rates maintained RH near 65% throughout the experiment, consistent with enhanced air exchange and moisture removal (Fig. 4C). Although lower exhaust flow improved particle transport, the associated increase in humidity was expected to impair aerosol stability and potentially affect animal welfare during long term studies. Based on these considerations, we selected an exhaust flow rate of 2.5 L/min for subsequent Mtb transmission experiments using the system 2B cage design. Together, these experiments defined an airflow regime that balanced aerosol transport with biological feasibility but did not explain why effective exposure differed across housing designs.

### Computational modeling explains particle retention differences across housing systems

To address this gap, we used computational fluid dynamics simulations to examine how airflow patterns and cage architecture influence particle retention and animal exposure across housing systems. Steady-state incompressible Navier-Stokes equations coupled to particle tracer advection-diffusion were solved to characterize airflow structure and aerosol transport within each system under both ambient conditions and BSL-3 negative-pressure laboratory environments (Supplemental Methods). These simulations enabled interpretation of experimental differences in particle retention and spatial distribution across housing systems. Full details of the computational framework, boundary conditions, and sensitivity analyses are provided in the Methods and Supplemental Methods.

We first defined the computational geometry and boundary conditions corresponding to the donor-recipient cage systems and surrounding laboratory environment (Fig. 5A-B). Simulations were performed for each system under both ambient pressure and BSL-3 negative-pressure conditions, incorporating tracer injection and exhaust locations matched to the experimental particle release and sampling setup. In the simulations, particle concentrations were evaluated within the naïve guinea pig breathing zone (BZ), defined as the cage volume below the animal’s typical breathing height of approximately 7 cm.

**Figure 5.**
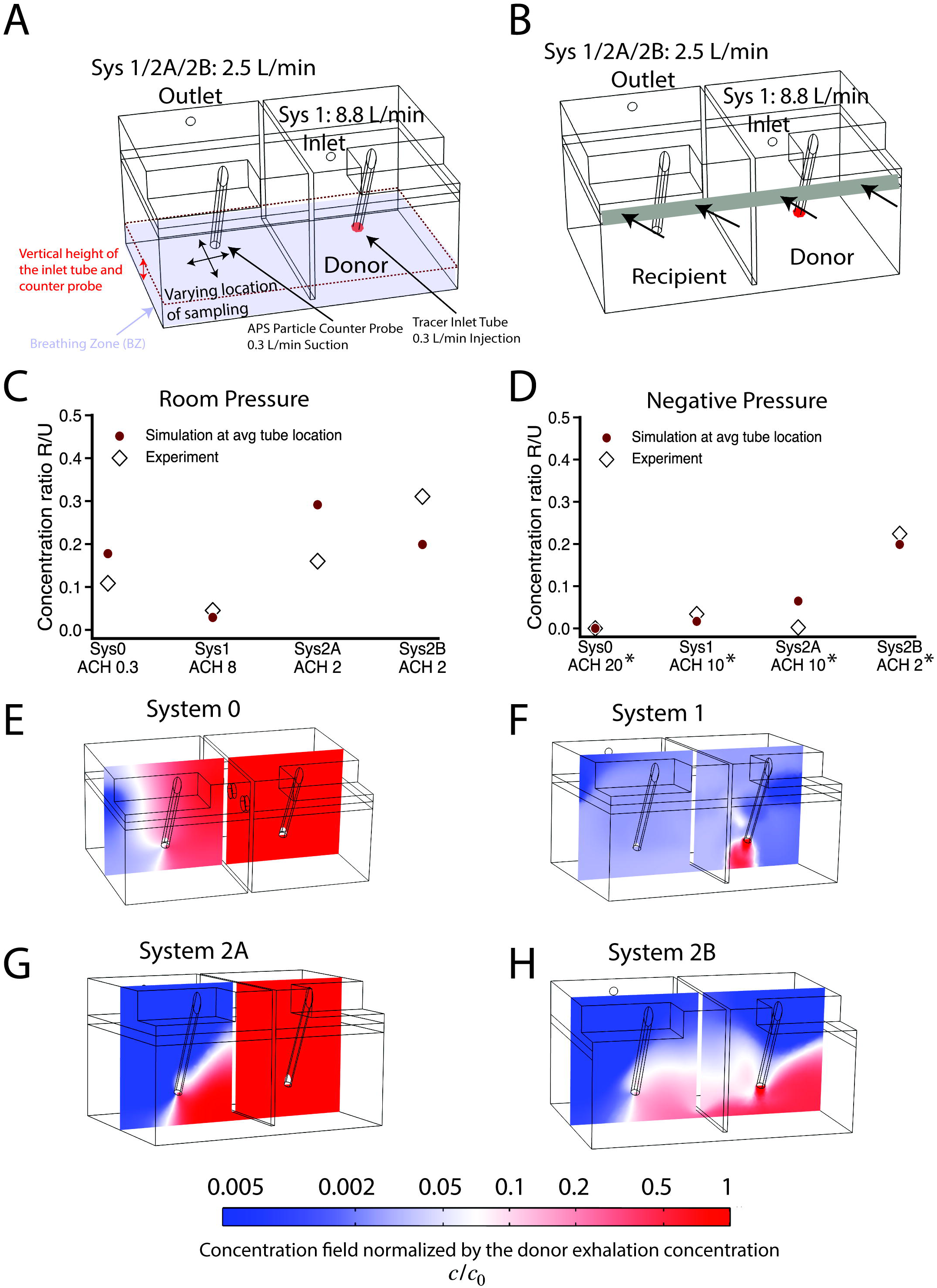
Particle tracer simulations, validated against experiments, reveal heterogeneous aerosol distributions and the influence of BSL-3 containment on particle retention. (A-B) Schematics of the donor-recipient cage system under ambient and negative-pressure surrounding laboratory conditions, showing airflow pathways and tracer injection and exhaust locations. The naïve guinea pig breathing zone (BZ) is defined as the cage volume below the animal’s typical breathing height of approximately 7 cm. (C-D) Comparison between computed and experimentally measured proxy for particle transport, quantified as the ratio (R/U) of tracer concentration in the recipient compartment (R) to upstream injected tracer concentration (U), shown for ambient (C) and negative-pressure *=negative pressure-induced effective ACH (D) laboratory conditions. Simulation values represent averages across four locations within the recipient-side BZ, capturing spatial variation in predicted particle concentrations; individual location-specific contributions are shown in Supplemental Fig. 2A-B. Experimentally measured R/U values are derived from single-location particle measurements and summed over particle size. Air-change rates per hour (ACH) for each configuration are indicated for ambient conditions. (E-H) Complementary animations for ambient and negative-pressure conditions are shown in Supplemental Movie 2 and 3

We quantified particle transport using the ratio of particle concentration in the recipient compartment (R) to that introduced upstream in the donor compartment (U). The computed R/U values were compared with the corresponding experimentally measured concentration ratios reported in Figs. 2B-I and 3B-I, summed over particle size. For the simulations, R/U is shown as the average across four locations within the recipient-side BZ (Fig. 5C-D), reflecting spatial variation in predicted particle concentrations (see below), with individual location-specific contributions shown in Supplemental Fig.2 A-B. Despite differences in sampling resolution, simulated and experimental R/U values tracked closely across systems and environmental conditions (Fig. 5C-D). In particular, simulations recapitulated the pronounced reduction in R/U observed experimentally under BSL-3 negative-pressure conditions for systems 0 and 2A and confirmed that system 1 remained particle-depleted due to its high ventilation rate.

To understand the spatial variation underlying these averaged R/U values, we next examined the simulated particle concentration fields across each housing system. These analyses revealed a clear breakdown of the well-mixed assumption, with strongly inhomogeneous aerosol distributions in most systems under both ambient and BSL-3 conditions (Fig. 5E-H; Supplemental Movies 2, 3). Spatial heterogeneity was most pronounced in system 2A, where sharp concentration gradients were observed within the recipient compartment (Fig. 5G). This heterogeneity is reflected in increased variability among the location-specific R/U values for system 2A and indicates that the inferred transmission proxy can be sensitive to measurement probe position in systems with non-uniform airflow patterns.

By contrast, system 1 exhibited relatively uniform but low particle concentrations due to rapid clearance, while system 2B maintained higher and more stable particle concentrations within the recipient-side BZ under both ambient and BSL-3 conditions (Fig. 5E-H, Supplemental Movies 2, 3). Importantly, these analyses showed that air changes per hour alone were insufficient to explain particle retention in this experimental context. Although systems 2A and 2B exhibited higher ventilation rates than system 0 under ambient conditions, localized airflow pathways and spatial heterogeneity dominated effective exposure within the recipient compartment.

Together, these simulations provide a physics-based interpretation of the particle tracer experiments and explain why only system 2B maintained effective aerosol retention under BSL-3 conditions, motivating its selection for subsequent transmission studies.

### Buoyancy from animal heat and respiration reshapes exposure patterns

Although particle tracer experiments and simulations captured key features of airflow-driven transport, these analyses initially treated particles as passive tracers in an isothermal flow. In living systems, however, animals introduce additional physical forces through body heat and respiratory exhalations that could plausibly reshape local airflow and aerosol transport. To account for these effects, we extended our simulations to incorporate buoyancy arising from guinea pig body heat and exhaled air, using physiologic parameters drawn from the literature, including animal size, body temperature, respiratory rate, tidal volume, and characteristic animal dimensions (*43–46*) (see Methods and Supplemental Methods for details of equations and parameters).

Guinea pigs generate continuous thermal plumes and episodic respiratory exhalations. A thermal plume is the gentle upward airflow produced by body heat, as warm air near the skin becomes buoyant and rises. In animals, this persistent flow can transport exhaled aerosols away from the body and interact with ambient ventilation. These host-derived flows interact with background ventilation and cage geometry in ways that might alter aerosol mixing, persistence, and access to the recipient-side breathing zone (BZ). We therefore used buoyancy-resolving simulations to assess whether host-generated forces could meaningfully influence effective exposure under BSL-3 housing conditions.

To quantify the biological implications of these buoyancy effects, we calculated a normalized, time-averaged particle concentration within the recipient guinea pig’s BZ, expressed relative to the concentration of particles emitted from the donor animal (𝒞; Fig. 6A; see Methods). This metric captures cumulative exposure experienced by the naïve animal under steady-state housing conditions and enables direct comparison across system designs without reliance on absolute particle counts.

**Figure 6.**
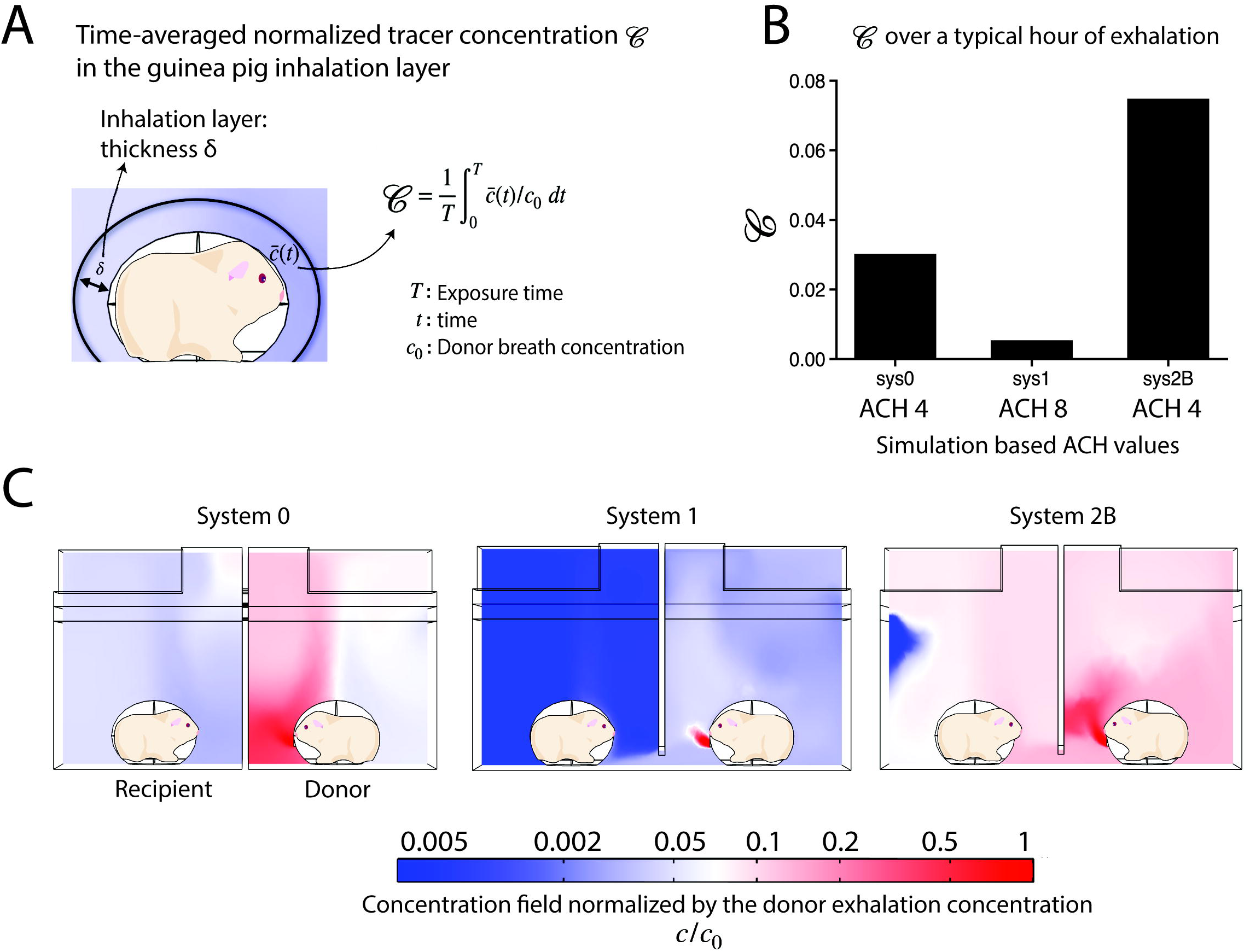
Buoyancy-resolving simulations quantify recipient exposure. (A) Definition of the recipient inhalation layer used to compute mean exposure (𝒞) (see Supplemental Methods). (B) Typical normalized time averaged exposure, 𝒞, of the recipient guinea pig for systems 0, 1 and 2B under negative pressure. (C) Representative mid-plane normalized tracer concentration field (c/*c*_0_), where *c*_0_ is the donor emission concentration (darkest red).

Using this exposure metric, simulations incorporating buoyancy predicted substantially higher recipient-side exposure in system 2B compared to earlier designs (Fig. 6B,C). Under otherwise equivalent conditions, system 2B produced approximately an order-of-magnitude greater normalized exposure than system 1, consistent with its enhanced particle retention. Visualization of the underlying flow fields revealed that buoyancy-driven upward mixing in system 2B promoted transfer of particles from donor to recipient compartments, whereas poorly sealed or highly ventilated systems diluted or redirected aerosols away from the recipient breathing zone (Fig. 6C; Supplemental Movie 5).

Enhanced buoyancy-induced mixing was further evident in mid-plane concentration slices of pathogen tracer distribution (Fig. 6C) and in the higher buoyancy-driven flow velocities observed across systems (Supplemental Fig. 3A-F and Supplemental Fig. 4A-F). Comparisons of simulations performed with and without buoyancy demonstrated that inclusion of animal body heat and respiratory exhalations substantially altered predicted airflow structure and particle transport pathways (Supplemental Movies 4, 5), highlighting the importance of host-derived forces in shaping exposure within enclosed transmission systems.

Together, these analyses indicate that airflow patterns imposed by cage design and ventilation interact with host-generated buoyancy to shape effective exposure in transmission systems. While airflow rate alone was insufficient to predict transmission potential, the combined effects of ventilation, sealing, and animal-derived buoyancy provide a mechanistic explanation for why system 2B uniquely supported sustained exposure. These insights informed selection of housing conditions for subsequent Mtb transmission experiments.

### Mtb transmission under optimized conditions

Having established airflow and environmental parameters that favored particle retention, we next assessed whether these optimized conditions supported Mtb transmission between guinea pigs. We infected guinea pigs with Mtb (Supplemental Fig. 5A-B) and housed them in the donor compartment of system 2B cages, while naïve guinea pigs were placed in the recipient compartment for a 17-week exposure period. Because guinea pigs develop progressive disease following infection, donor animals were replaced every 6 weeks to maintain a consistent infectious source.

We monitored transmission using monthly tuberculin skin testing (TST) with a two-stage protocol (*47–49*) in which animals were tested twice with a 7-day interval between doses (Fig. 7A). Donor animals developed robust TST responses by four weeks post-infection (mean 10 mm; range 6-16 mm), consistent with active infection (Fig. 7B). To define a threshold for positive conversion in recipients, we performed repeated TSTs in unexposed control animals and found that none exceeded 5.5 mm (Supplemental Fig. 5D). Using this baseline, together with TST responses from infected donors (Fig. 7B) and uninfected naïve animals (Supplemental Fig. 5E), we established a cutoff of 6.5 mm to indicate infection. Within one month of exposure, one recipient exceeded this threshold, and by week 17, nine of twelve recipient animals demonstrated positive TST conversion (Fig. 7B; Supplemental Fig. 6).

**Figure 7:**
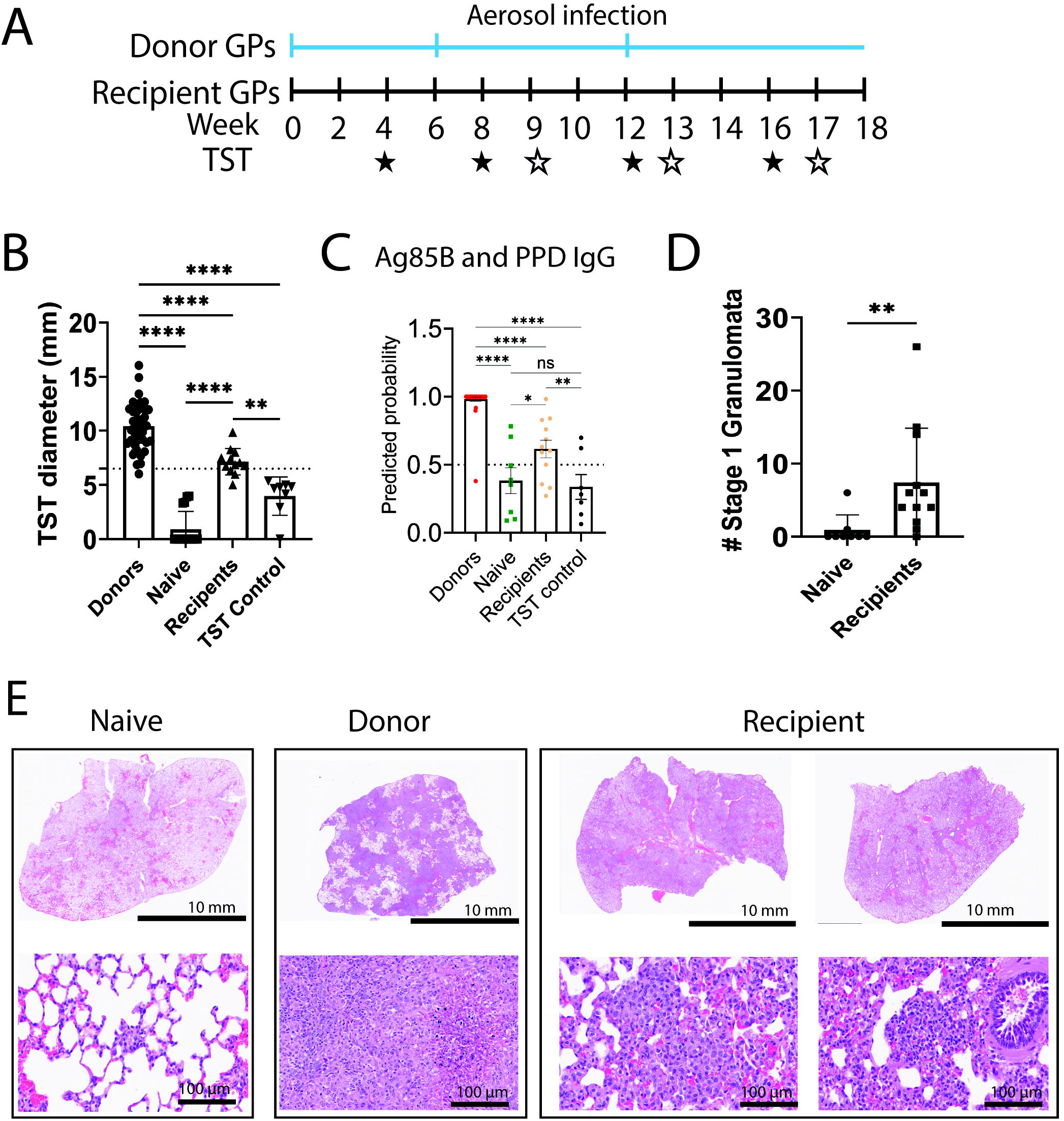
Transmission of Mtb in an optimized cage system. (A) Experimental timeline for 17-week transmission study. Blue ticks indicate when new donor groups were added to the cages. Monthly TSTs using a two-step strategy are noted as filled stars (first dose) and open stars (second dose). (B) TST measurements for recipient animals throughout the exposure period. Any responses above 6.5 (dashed line) are considered positive. (C) Predicted probability of Mtb exposure based on serum antigen 85B and PPD IgG levels. Values above 0.5 (dashed line) are predicted positive in the donor, naïve, recipient and TST control groups. (D) Quantification of stage 1 granulomata in naïve and recipient lung sections. (E) Representative hematoxylin and eosin staining of lung section from naïve, donor and recipient groups. Inflammatory collection (naïve) and stage 1 granuloma (recipient) at 20x. p-values calculated by one-way ANOVA for multiple comparisons or Mann-Whitney for comparison between two groups. ns=not significant, *=p<0.05, **= p<0.01, ***=p<0.001, ****=p<0.0001.

Because Mtb exposure also elicits humoral immune responses (*50–55*), we next quantified serum antibody responses to Mtb antigens using a bead-based multiplex assay (Supplemental Fig. 7A) (*55*). Antigens included complex mixtures such as PPD and defined immunodominant proteins such as antigen 85B. Area-under-the-curve (AUC) values were calculated from dilution series, and receiver operating characteristic (ROC) analyses were performed for individual and combined antigen-specific IgG responses (Supplemental Fig. 7B-G). Infected donor animals exhibited strong IgG responses, whereas unexposed controls remained negative. Using a predictive probability cutoff of 0.5 for the combined IgG response to antigen 85B and PPD (Supplemental Fig. 7H), eight of twelve recipient animals displayed serologic responses consistent with Mtb exposure (Fig. 7C).

We next examined lung tissues from recipient guinea pigs by blinded histopathology analysis. Compared to unexposed controls, exposed recipients exhibited increased pulmonary inflammation characterized by stage 1 granulomas, intra-alveolar hemorrhage, and small foci of epithelioid macrophages (Fig. 7D-E, Supplemental Fig. 5F-G). Although recipient animals did not exhibit weight loss (Supplemental Fig. 5C) and no culturable Mtb was recovered at necropsy, the concordance of TST conversion, antigen-specific serologic responses, and histopathologic changes indicates that recipient animals were exposed to and mounted immune responses to Mtb.

Together, these findings demonstrate that the optimized system 2B configuration supports natural airborne transmission of Mtb between infected and naïve guinea pigs. By integrating controlled airflow, cage sealing, and structural refinement, this system enables sufficient aerosol retention to permit biological transmission under BSL-3 containment, providing a reproducible experimental platform for studying host, microbial, and environmental determinants of airborne tuberculosis spread.

## DISCUSSION

Our current understanding of TB transmission arises largely from epidemiologic studies of Mtb spread within human populations or from analysis of exhaled aerosols (*24, 25, 28, 56–58*). Experimental validation of these observations has been constrained by the absence of modern animal-to-animal transmission systems that operate under contemporary biosafety conditions. Here, we establish a guinea pig model that supports natural airborne transmission of Mtb under BSL-3 containment, enabling direct experimental interrogation of the biological determinants of transmission.

Using iterative cage design and controlled transmission experiments, we identified airflow conditions that permit sufficient aerosol persistence to support biological exposure. Transmission required retention of particles in the 1-5 µm aerodynamic diameter range capable of carrying Mtb (*24, 25, 28, 57*), while remaining compatible with animal welfare and biosafety constraints. Under these optimized conditions, recipient guinea pigs mounted antigen-specific immune responses and developed pulmonary inflammation consistent with early infection, demonstrating that natural airborne transmission can be reconstituted experimentally.

Importantly, without tight seal or flow pattern control, both extremes of ventilation suppressed transmission. Static environments, despite low nominal air-change rates, permitted particle loss through uncontrolled leakage and buoyancy-driven escape, limiting delivery of aerosols to the recipient breathing zone. Conversely, excessive unidirectional ventilation rapidly depleted aerosolized particles before effective exposure could occur. These findings demonstrate that ventilation rate alone is insufficient to predict transmission risk and highlight the importance of local airflow patterns, enclosure integrity, and host-generated forces in determining biological exposure.

The influence of overall ventilation on infection risk has long been recognized from the classic studies of Riley and Wells to modern public health strategies for airborne disease control. Renewed attention to ventilation during the COVID-19 pandemic emphasized air exchange as a tool to mitigate transmission in real-world settings (*59*). Our results provide experimental confirmation of this principle, showing that sufficiently high ventilation suppresses transmission even in close-proximity animal housing. At the same time, they underscore the limitations of the well-mixed air assumption and demonstrate that local flow topology and enclosure design critically shape exposure under biosafety-mandated ventilation conditions. In addition, our results provide experimental confirmation that airborne TB transmission can be mitigated with sufficient replenishment of fresh air (*48, 60–63*), as illustrated by our experimental outcomes showing absence of animal-to-animal transmission in systems 0 and 1 under BSL3-containment conditions.

By integrating particle tracer experiments with computational analyses, we show that airflow heterogeneity and buoyancy generated by live animals substantially alter aerosol transport within confined housing systems. These physical insights help explain why only specific cage designs supported sustained exposure and transmission, despite similar average ventilation metrics. Accounting for these effects provides a mechanistic explanation for the historical difficulty of establishing animal-to-animal transmission models in modern biosafety facilities.

The ability to reproduce natural transmission has important implications for tuberculosis biology. Few experimental systems currently allow direct testing of how bacterial, host, and environmental factors interact to shape the transmission cascade. Existing approaches rely on artificial aerosolization of Mtb to naïve animals (*64*), human-to-animal exposure systems (*23, 65*), or alternative animal models with uncertain relevance to human disease (*15, 16*). The system described here enables transmission to be studied directly between infected and naïve animals in a well-established model of tuberculosis pathogenesis. For example, this platform can be adapted to investigate microbial determinants of transmission such as strain lineage (*66–72*), lipid composition (*73–75*), genetic background (*76*) or specific Mtb genes (*37*). It also provides an opportunity to examine how donor physiology influences aerosol generation and infection probability.

The relationship between respiratory behavior and transmission remains a particularly important gap in our understanding of Mtb transmission biology. Although tidal breathing alone can generate infectious bioaerosols (*32, 77–79*), coughing produces larger quantities of particles and propels them over longer distances (*28, 33, 39, 59, 80, 81*). Our prior work identified Mtb cell wall glycolipids sufficient to induce cough in naïve guinea pigs (*73, 74*), and guinea pigs infected with Mtb strains lacking both lipids did not cough, despite severe pulmonary pathology (*74*). Applying the current transmission model to such strains will allow direct testing of how pathogen-encoded cough-inducing factors influence transmission efficiency. In the present simulations, breathing and coughing were assigned equivalent emitted pathogen concentrations to isolate the role of airflow and host-generated forces in shaping exposure, independent of variation in bacterial source. Although a single cough is expected to generate a larger number of particles than a single breath, breathing occurs orders of magnitude more frequently over time, making both processes biologically relevant contributors to cumulative exposure. Our controlled framework can be extended in future studies to explicitly vary source strength and respiratory behavior to define how cough frequency and intensity interact with airflow to influence transmission efficiency.

This study has limitations. Although recipient animals showed consistent immunologic and histologic evidence of exposure, culturable Mtb was not recovered from lungs at necropsy. Similar findings have been reported in human-to-guinea pig transmission studies, where latent infection was revealed only after immunosuppression (*23*). In addition, replacement of moribund donor animals precluded correlation of donor bacterial burden or cough frequency with transmission outcomes. Longer exposure durations, recovery phases, or transient immunosuppression may increase bacterial detection; future studies will allow donor disease burden to be correlated with transmission efficiency. Our modeling approach employed a continuum particle tracer rather than explicitly tracking discrete particles of varying size and deposition behavior. While this simplification limits quantitative inference about individual particle fate, it preserves the dominant airflow structures governing aerosol transport within the system. Accordingly, our conclusions primarily reflect relative exposure patterns and transmission-relevant airflow dynamics rather than absolute particle deposition.

In summary, this work revives a classical model of airborne tuberculosis transmission and adapts it for modern biosafety conditions. By defining airflow regimes that permit biologically effective exposure, we establish a reproducible experimental system that supports natural Mtb transmission between animals. This platform enables mechanistic investigation of the microbial, host, and environmental factors that govern airborne spread and provides a foundation for developing transmission-blocking interventions.

## MATERIALS AND METHODS

### Animal studies

Protocols for guinea pig infection and transmission studies were reviewed and approved by the Institutional Animal Care and Use Committee at the University of Texas Southwestern Medical Center (Protocol # 2018-102280-USDA).

### Particle transmission

Aerosol particles were generated using a custom-built particle generation system to model transport of respirable airborne particles under defined airflow conditions. A syringe pump (Braintree Scientific, MA, USA) delivered EMEM (Suppl. Fig 1B) suspension solution at a flow rate of 0.4 mL/min. Compressed air at 60 psi atomized and dried the aerosols, with a total flow of 21.5 L/min. Relative humidity along the tubing was monitored with a probe (Ahlborn, Germany) and maintained at 48 ± 2%. Continuous aerosol loading into the cage was controlled at 0.3 L/min, while excess flow was vented using a vacuum pump (Gast Manufacturing, MI, USA) regulated by a mass flow controller (MFC; Aalborg Instruments and Controls, NY, USA).

Four case systems were evaluated: System 0, System 1, System 2A, and System 2B. Each consisted of donor and recipient compartments separated by a divider, with 0.3 L/min entering the donor side. Flow conditions varied among tests as summarized in Table 1. Compressed supply air (8.8 L/min) could be directed into the donor side, and exhaust air (2.5 L/min) was regulated using a custom-built laminar flow element connected to the facility vacuum.

**Table 1:**
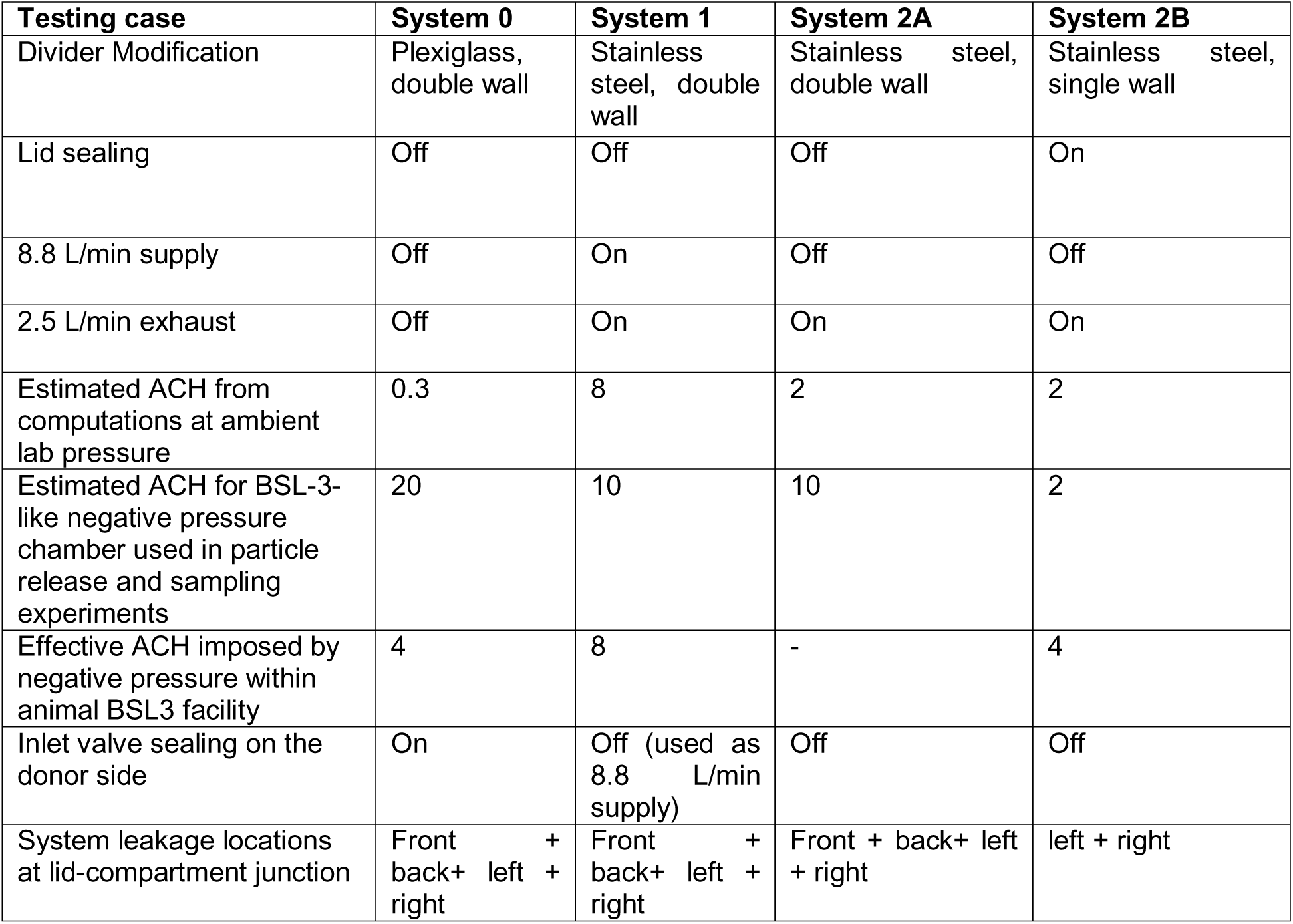
Structural and airflow characteristics of guinea pig transmission housing systems.

Aerosol particle size distributions were measured on both donor and recipient sides using an Aerodynamic Particle Sizer (APS; Model 3321, TSI, MN, USA). The APS drew 0.3 L/min directly from each sampling site, while dilution air was added through compressed air controlled by a MFC to achieve the required 5 L/min sampling flow. Before aerosol introduction, clean air was flushed through the cage for 30 min to confirm negligible background particle concentrations. Aerosol particles were then loaded for approximately 1 h to reach steady-state size distributions (Supplemental Fig. 1A). After loading, three 20-second samples were collected per minute, alternating between donor and recipient sides. The particle size boundary was set to 0.8-7µm. Sampling was repeated every 5 min over a 25-min period under both ambient and negative-pressure conditions.

To reproduce biosafety level 3 negative-pressure environments, a custom chamber (24″ × 24″ × 48″) was constructed. An inline duct fan (TerraBloom LLC, CA, USA) regulated airflow, and chamber pressure was monitored continuously using a differential gauge (Dwyer Instruments, Inc., IN, USA), maintained at −0.03 or −0.06 inches of water.

### Mycobacterium smegmatis particle transmission

*Mycobacterium smegmatis* cultures were grown in Middlebrook 7H9 broth (Sigma #M0178) supplemented with 10% OADC, 0.5% glycerol, and 0.05% Tween-80 at 37 °C with shaking (100 rpm). Cultures were diluted to 1 × 10 CFU/mL in PBS and aerosolized using a Blaustein Atomizing Module (CH Technologies). The atomizer was placed in the donor compartment of the transmission cage, and aerosols were generated for 1 min. After a 2-min delay, air from the recipient compartment was sampled for 30 s using a button sampler (SKC, PA, USA) with a gelatin filter at 10 L/min. The filter was dissolved in PBS and plated on Middlebrook 7H10 agar supplemented with OADC and glycerol. Colonies were counted after overnight incubation to quantify recovered bacteria. This procedure was repeated at exhaust flow rates of 2.5, 1.5, and 0.15 L/min.

For transmission of *M. smegmatis* to guinea pigs, the atomizer was placed in the donor compartment containing 1 × 10^5^ CFU/mL of *M. smegmatis* in PBS. The recipient compartment contained 2 guinea pigs. Aerosols were generated for 1 min and allowed to pass to the recipient compartment and inhaled by the recipient animals for 15 minutes. Following the 15-minute period, animals were sacrificed and lungs were homogenized in PBS and plated on four 7H10 agar plates per animal to quantify recovered bacteria. This process was repeated for a total of four guinea pigs.

### Guinea pig aerosol infection

Donor guinea pigs were infected with *M. tuberculosis* Erdman in three groups on a 6-7 week cycle. Bacteria were cultured in Middlebrook 7H9 broth supplemented with 10% OADC, 0.5% glycerol, and 0.05% Tween-80 at 37 °C using a roller apparatus. Cultures were washed three times with PBS, then centrifuged at 500 rpm for 5 min to remove large clumps. The supernatant was sonicated at 90% amplitude for 15-s pulses (Branson Ultrasonics, CT, USA) to generate a single-cell suspension and diluted to 3 × 10 CFU/mL in PBS. The inoculum was aerosolized using a Glas-Col inhalation exposure system (Glas-Col, IN, USA). Two animals per group were sacrificed immediately post-exposure for day-0 CFU quantification by lung plating on 7H11 agar. All animals were weighed weekly, and donors were euthanized seven weeks post-infection.

### Transmission experiments and cage modifications

For transmission experiments, male Hartley guinea pigs (Charles River) were used. For system 0, we used a polycarbonate static cage (Tecniplast # 2000P00SU), with a raised lid (Tecniplast #2000P224) covered by a filter top (Tecniplast #U1144B420RBOX-300) with a custom designed double walled plexiglass divider perforated with holes for particle movement between compartments. For the transmission experiment, cages were maintained on a stainless-steel rack in our ABSL3. For systems 1 and 2B, we housed animals in an Allentown 1800 15 cage rack (Allentown, NJ, PA). The standard cage bottom (Allentown #315351-1) and external bottle top (Allentown #230347) were used with the following modifications. For systems 1, 2A and 2B, the donor-side exhaust port (right) and recipient-side supply ports (left) were sealed with rubber caps. While for system 1 the supply air was “on”, for systems 2A and 2B, the supply hose was disconnected from the main air inlet of the rack. For system 2B, a custom stainless-steel cage divider with 102 ventilation holes was fabricated locally to separate donor and recipient animals within each cage. To improve sealing, the underside of the lid was covered with aluminum tape, a 1.6 mm rubber gasket was applied along the lid-base junction, and four 2-inch spring clamps secured the lid once cages were positioned on the rack.

### Tuberculin skin test

To determine Mtb exposure, guinea pigs were tested every four weeks using a two-step TST protocol. Animals were anesthetized under 3% isoflurane, and the right or left flank was shaved and cleaned with 70% ethanol. Tuberculin purified protein derivative (PPD; 100 µL of 5 TU, McGuff Medical Products #001079) was injected intradermally. Induration was measured at 24 h and 48 h post-injection. The procedure was repeated one week later on the opposite flank. Control animals received a single dose of tuberculin at baseline or the same two-step PPD dosing scheme as exposed animals to control for effects of repeated testing.

### Serologic analysis

Antigen-specific IgG levels were quantified as previously described with minor modifications (*55, 82, 83*). Carboxylated microsphere beads (Luminex Corp., Austin, TX, USA) were coupled to protein antigens via covalent NHS-ester linkages using 1-ethyl-3-[3-dimethylaminopropyl] carbodiimide hydrochloride (EDC) and *N*-hydroxysulfosuccinimide (NHS) (Thermo Fisher Scientific, Waltham, MA, USA), following the manufacturer’s instructions. MagPlex® Avidin microspheres (Luminex) were conjugated to biotinylated sulfolipid-1 (SL-1) using the manufacturer’s protocol.

The following antigens were used: Ag85A (BEI Resources NR-14871), Ag85B (NR-14870), ESAT-6 (NR-49424), CFP-10 (NR-49425), purified protein derivative (PPD; Statens Serum Institute, Copenhagen, Denmark), *M. tuberculosis* H37Rv whole-cell lysate (NR-14822), and *M. tuberculosis* H37Rv cell wall fraction (NR-14828).

Antigen-coupled microspheres (1,000 beads per well) were incubated with diluted guinea pig serum (1:50, 1:250, and 1:1,250) in 96-well Bio-Plex Pro flat-bottom plates (Bio-Rad Laboratories, Hercules, CA, USA) at 4 °C for 16 h. After washing to remove unbound antibodies, antigen-specific IgG was detected using phycoerythrin (PE)-conjugated goat anti-guinea pig IgG (Novus Biologicals, NBP1-74874PE). Beads were incubated with the secondary antibody for 2 h at room temperature, washed with PBS containing 0.05% Tween-20, and resuspended in PBS. Fluorescence intensity was acquired using a MAGPIX® instrument running xPONENT 4.2 software (Luminex).

Relative antigen-specific IgG levels were quantified as the area under the curve (AUC) calculated from the three serial serum dilutions for each sample.

Receiver operating characteristic (ROC) curves were generated using GraphPad Prism version 10.5 (GraphPad Software, San Diego, CA, USA). Sensitivity and specificity at selected cutoff points, along with 95% confidence intervals, were calculated using logistic regression.

### Histologic analysis

At study completion, animals were euthanized, and lungs were collected in 10% neutral-buffered formalin for histopathologic analysis. Fixed tissues were transferred to 70% ethanol and processed by the Histo Pathology Core Facility at UT Southwestern. Paraffin-embedded blocks were sectioned at 5 µm and stained with hematoxylin and eosin. Slides were scanned using a NanoZoomer S60 digital slide scanner, randomized, and blinded prior to evaluation by a pathologist. The number and stage of granulomas were recorded, and pulmonary inflammation was described for each animal. After unblinding, data were compared between naïve and exposed animals using a Mann-Whitney test (GraphPad Prism v10).

### Computational modeling of airflow and particle transport within transmission housing systems

To interpret experimentally observed differences in particle retention and exposure across housing designs, we used computational fluid dynamics simulations to model airflow and tracer particle transport within the transmission cages. Simulations were performed under both ambient laboratory conditions and negative-pressure BSL-3 environments, using cage geometries and airflow boundary conditions matched to experimental configurations.

Airflow was modeled as incompressible, steady-state flow, and aerosol particles were treated as passive tracers subject to advection and diffusion. To account for host-generated effects present in occupied cages, additional simulations incorporated buoyancy arising from animal body heat and respiratory exhalations using physiologic parameters drawn from the literature, including guinea pig size, body temperature, respiratory rate, tidal volume, and characteristic body dimensions.

Full governing equations, numerical methods, boundary conditions, parameter values, and sensitivity analyses are provided in the Supplemental Methods.

### Metric of relative exposure used to compare systems

To enable biologically meaningful comparison of exposure across housing designs, we defined a normalized, time-averaged exposure metric, 𝒞, representing the average concentration of particles within the recipient guinea pig breathing zone relative to the concentration emitted by the donor animal. The breathing zone was defined as the region from which inhaled air is predominantly drawn during normal respiration.

This metric captures cumulative inhalation exposure over time and allows comparison across systems independent of absolute particle emission rates. The mathematical definition of 𝒞, spatial averaging procedures, and parameter choices are provided in the Supplemental Methods.

## ACKNOWLEDGMENTS

The authors would like to thank Cody Ruhl for preliminary experiments, and all members of the Shiloh lab for their support and constructive feedback. We also thank members of the Animal Resources Center at UT Southwestern for providing support with animal husbandry and the engineering team at Allentown, Inc. for their feedback throughout this project.

## AUTHOR CONTRIBUTIONS

Conceptualization, K.F.N, L.B. and M.U.S. Formal Analysis, K.F.N, Y.G., Y.K., D.S., P.L., S.W., A.M., B.M.E., L.L., H.O., L.B., and M.U.S. Funding Acquisition, L.B. and M.U.S. Investigation, K.F.N, Y.G., Y.K., D.S., P.L., S.W., B.R.S.D., V.A.E., A.E.M., B.M.E., L.L., and L.B. Project Administration, K.F.N and M.U.S.Resources, H.O., M.U.S. Validation, K.F.N, Y.G., D.S., P.L., A.E.M., Y.K. Supervision, L.L.L., H.O., L.B. and M.U.S. Writing-original draft, K.F.N, Y.G., Y.K., H.O., L.B., and M.U.S. Writing-Review and Editing, all authors.

