## Supplemental Methods and Figures for "Airflow Constraints Limit Natural Airborne Transmission of Tuberculosis"

### NUMERICAL SIMULATION METHODOLOGY

Simulations were performed using COMSOL Multiphysics to model airflow and tracer transport within the guinea pig transmission housing systems. The computational domain was constructed to match the physical dimensions of the Allentown NexGen™ 1800 cage system used experimentally, including donor and recipient compartments, separators, and leakage pathways specific to each housing configuration (systems 0, 1, 2A, and 2B).

#### 1.Tracer particle-only steady-state simulations (Systems 0, 1, 2A, and 2B)

We used the tracer particle-based exposure experiments discussed in Figs. 2 and 3 (main text) to 1) benchmark our computations; and 2) furnish the underlying physical mechanisms governing the experimental sampling measurements and their variability. Airflow and tracer transport were simulated under steady-state conditions using COMSOL Multiphysics, coupling laminar airflow with passive tracer transport.

#### 1.1 Governing equations

The flow continuity, momentum balance, and the tracer particle advection-diffusion equations are coupled and solved for the fluid velocity, $\boldsymbol{u}$, and tracer concentration $c$, fields. They read:

$$\nabla \cdot \boldsymbol{u} = 0, \tag{1.1}$$

$$\rho(\boldsymbol{u} \cdot \nabla)\mathbf{u} = -\nabla p + \mu \nabla^2 \boldsymbol{u}, \tag{1.2}$$

$$\nabla \cdot (-D \nabla c + \boldsymbol{u} c) = 0. \tag{1.3}$$

The parameters used above are given in **Table S1**.

| Parameter | Symbol | Value | Applicable case |
|---|---|---|---|
| **Air density** | $\rho$ | 1.20 kg $m^{-3}$ | Particle, GP |

| | | | |
|---|---|---|---|
| **Dynamic viscosity** | $\mu$ | $1.81 \times 10^{-5}$ Pa·s | Particle, GP |
| **Ambient pressure** | $p$ | 1 atm | Particle, GP |
| **Ambient temperature** | $T_0$ | 293.15 K (20 °C) | Particle, GP |
| **Scalar diffusivity** | $D$ | $1 \times 10^{-6}$ m² s$^{-1}$ | Particle |
| **Scalar diffusivity (exhaled tracer)** | $D$ | $2 \times 10^{-5}$ m² s$^{-1}$ | GP |
| **Thermal expansion coefficient** | $\beta$ | $3.4 \times 10^{-3}$ K$^{-1}$ | GP |
| **Specific heat capacity** | $c_p$ | 1005 J kg$^{-1}$ K$^{-1}$ | GP |
| **Thermal conductivity** | $k$ | 0.026 W m$^{-1}$ K$^{-1}$ | GP |

**Table S1 . Summary of physical and tracer pathogen transport parameters used in particle-tracer computations.**

#### 1.2 Boundary conditions

For all systems (systems 0, 1, 2A, and 2B), no-slip and non-penetration boundary conditions were applied for both the air and the tracer particles at the walls $\Gamma_{\text{wall}}$: $\boldsymbol{u} = \mathbf{0}$ and $(-D\nabla c) \cdot \boldsymbol{n} = 0$, where $\mathbf{n}$ is the normal to the walls, $\Gamma_{\text{wall}}$.

For systems 1, 2A, and 2B, the recipient side was exposed to donor emissions only via tracer transport through the bottom gap between the separators, whereas for system 0, transport occurred exclusively through the open holes in the upper part of the separator (Fig. 5 A-B).

Flow-driven boundary conditions were imposed assuming fully developed flow profiles with specified volumetric flow rate,

$$\int_{\Gamma_{\text{in}}} \mathbf{u} \cdot \mathbf{n} \, dA = Q,$$

with $\mathbf{u} \cdot \mathbf{t} = 0$, where $\mathbf{t}$ denotes the unit tangent vector along the boundary.

For all systems, the outward flow rate at the recipient side was fixed at 2.5 L/min. The back-wall opening acted as an air inlet for system 1, where a fully developed flow profile was imposed at 8.8 L/min, while for systems 2A and 2B this opening was treated as an open boundary (Fig. 5A-B).

Consistent with experimental conditions discussed in main text Figs. 2 and 3, tracer particles were injected at a rate of 0.3 L/min via tubing introduced in the compartments in the experiments (Fig. 5 and 6) to approximate the breathing zone of the guinea pigs.

**Leakage modeling**

Leakage strips, shown in Supplemental Fig. 8 A-B, were located at the junction between the cage top and bottom (external bottle cover interface). These regions were modeled as open boundaries for systems 0, 1, and 2A.

For tracer particle-only experiments in system 2B, the front and back leakage pathways were mechanically sealed with tight clamps. The side strips (left and right) were partially sealed using rubber gaskets that reduce but do not eliminate leakage. Accordingly, in the computational model we sealed the front and back strips and retained reduced side leakage strips modeled as resistive “grille” boundaries and at the same locations as in systems 0, 1, and 2A.

These grille boundaries were implemented as pressure-loss boundary conditions acting as lumped hydraulic resistances, commonly used to capture resistive interfaces with unresolved micro-geometry (e.g., ventilation grilles or perforated panels) where explicit meshing would be computationally prohibitive (*84, 85*).

In COMSOL, the grille boundary condition was implemented as a modified traction condition on the leakage strip $\Gamma_{\text{strip}}$,

$$\mathbf{n}^{\mathrm{T}}(-p\,\mathbf{I}+\mathbf{K})\mathbf{n}=-p_0, \qquad \mathbf{u}\cdot\mathbf{t}=0,$$

where $\mathbf{n}$ and $\mathbf{t}$ denote the outward unit normal and tangential vectors, $\mathbf{u}$ is the velocity vector, $p$ is the local fluid pressure, $\mathbf{I}$ is the identity tensor, $\mathbf{K}$ is the viscous stress tensor, and the superscript $(\cdot)^{\mathrm{T}}$ denotes the transpose operator.

The effective boundary pressure $p_0$ is constrained by $p_0 \geq p_{\mathrm{input}} - \Delta p_c$, with $p_{\mathrm{input}}$ the ambient reference pressure and $\Delta p_c$ a pressure-loss term determined by the *linear loss coefficient* (LLC). For a linear grille model, the pressure loss scales with the volumetric flow rate through leakage boundaries,

$$\Delta p_c \sim LLC\,Q_{\mathrm{strip}}, \qquad \text{with} \quad Q_{\mathrm{strip}}=\int_{\Gamma_{\mathrm{strip}}}\mathbf{u}\cdot\mathbf{n}\,dA$$

reflecting the inflow through the leakage strip. This formulation acts as a lumped hydraulic resistance, that is, for a given driving pressure environment, the leakage flow rate scales inversely with the loss coefficient,

$$Q_{\mathrm{strip}} \sim \frac{\Delta p_{\mathrm{drive}}}{\mathrm{LLC}}.$$

Thus, introducing a finite $LLC$ suppresses leakage without enforcing a perfectly sealed boundary. To interpret this resistance geometrically, note that to leading order, leakage flow, under similar pressure conditions, scales with an effective opening area $A_{\mathrm{eff}}$. Comparing a resistive leakage strip with an otherwise identical open strip yields the scaling relation

$$\frac{A_{\mathrm{eff}}}{A_0} \sim \frac{Q_{\mathrm{strip}}}{Q_{\mathrm{open}}},$$

where $A_0$ and $Q_{\mathrm{open}}$ denote the nominal strip area and the corresponding open-boundary leakage. Thus, in our case, the grille model corresponded to an effectively thinner leakage pathway relative to systems 0, 1, and 2A. For the present tracer particle-only simulations, we used an LLC value of 4 kg $m^{-4}$ $s^{-1}$ which produced an order-unity reduction in leakage flow relative to an open strip. Interpreted through the scaling relation above, this corresponded to an effective reduction in leakage thickness from nominal height of ≈ 2 cm to a value in the range of ≈ 5 mm to 1 cm, depending on the surrounding pressure environment.

**Tracer transport boundary conditions**

The tracer inflow concentration at the injection inlet is fixed at a normalized reference value $c_0 = 1\,mol/m^3$. Thereby, tracer concentration is normalized per unit volumetric inflow rate, to enable ease of concentration field comparison across systems.

The back-wall opening in system 1 injects fresh air and therefore assigned a particle concentration $c = 0$. Tracer outflow is permitted through all open boundaries, including leakage strips and back-wall openings when present. Incoming air through these boundaries is free of tracer particles.

Thus, leakage strips and back-wall openings are open boundary conditions with respect to tracer transport, such that $-\mathbf{n} \cdot \mathbf{J} = 0$ if $\mathbf{n} \cdot \mathbf{u} \geq 0$, corresponding to purely advective outflow and $c = 0$ if $\mathbf{n} \cdot \mathbf{u} < 0$, ensuring that any inflow through these boundaries consists of tracer-free air where $\mathbf{J}$ denotes the tracer flux.

**Mesh convergence**

To evaluate numerical robustness, we conducted a mesh-convergence study (*86, 87*) to ensure independence of the reported fields from grid resolution, ***h***; with details shown in Table S2. Five successively refined meshes were tested, parameterized by refinement factor ***h***, where each refinement halves ***h.***

| Mesh | Mesh refinement factor: h | Recipient-side concentration | % change from previous mesh grid resolution | % diff. from finest | Concentration at the probe location | % change from previous | % diff. from finest |
|---|---|---|---|---|---|---|---|
| 1 | 1 | 0.103 | - | 15.6% | 0.277 | - | 36.4% |
| 2 | 0.5 | 0.096 | -7.3% | 7.3% | 0.236 | -14.7% | 16.3% |
| 3 | 0.25 | 0.092 | -3.9% | 3.1% | 0.218 | -7.6% | 7.5% |
| 4 | 0.125 | 0.090 | -1.8% | 1.2% | 0.208 | -4.5% | 2.6% |
| 5 | 0.0625 | 0.089 | -1.2% | - | 0.203 | -2.5% | - |

**Table S2 . Mesh-convergence study** Mesh-convergence over successively refined meshes parameterized with the refinement factor h, where h=1 corresponds to Mesh 1 and each subsequent mesh is obtained by halving h. The minimum element size in the bulk domain was set to 2.5h mm, while the leak-strip, inlet-outlet, and probe regions used a finer minimum element size of 2.0h mm with a maximum element size of 5h mm. The maximum element growth rate was fixed at 1.2, and a curvature factor of 0.1 was applied uniformly across all meshes to ensure consistent refinement behaviour. To resolve sharp edges near the lower cage openings, the resolution of narrow regions was set to 0.9, and a corner-refinement element-size factor of 0.07 was maintained for all meshes. Two metrics are reported: the recipient-side (volume-averaged) concentration and the local probe concentration, obtained as an average concentration over the probe area of radius 1 cm. Percentage change relative to the previous mesh and percentage differences relative to the finest mesh (Mesh 5) are reported to assess convergence. Mesh 4 (h = 0.125) is used in this study.

Two scalar concentration metrics were evaluated:

1. the recipient-side volume-averaged tracer concentration, and

2. the local probe concentration, obtained by averaging tracer concentration over a fixed probe area of radius 1 cm.

The recipient-side concentration converged rapidly with mesh refinement, with differences below 1.2% between the two finest meshes. The probe concentration converged more slowly, as expected for a local metric, while exhibiting monotonic near first-order convergence, with a remaining difference of 2.6% between the two finest meshes. An example of coarse refinement and finer refinement is shown in Supplemental Fig. 8C-E. We use the finer refinement in all the computations discussed in the main text.

### 2. Guinea pig modeling: breathing and coughing exhalations

#### 2.1 Governing equations

For simulations involving guinea pig occupants, we coupled the Heat Transfer in Fluids module with the Laminar Flow and Transport of Dilute Species modules using COMSOL's Multiphysics framework. The Boussinesq approximation (*88, 89*), was used to account for buoyancy arising from temperature variations, while maintaining the assumption of incompressible flow. We solved the continuity and momentum equations for the velocity field $\mathbf{u}$, the energy equation for temperature $T$, and the advection-diffusion equation for tracer concentration $c$:

$$\nabla \cdot \mathbf{u} = 0,$$

$$\rho(\mathbf{u} \cdot \nabla)\mathbf{u} = -\nabla p + \mu \nabla^2 \mathbf{u} + \rho \mathbf{g}\, \beta (T - T_0),$$

$$\rho c_p (\mathbf{u} \cdot \nabla T) = k \nabla^2 T,$$

$$\nabla \cdot (-D \nabla c + \mathbf{u} c) = 0,$$

Where the parameters are defined in Table S1 (applicable case: GP). The scalar diffusivity of the exhaled tracer in air from the donor guinea pig mouth was consistent with molecular diffusion coefficients of gases such as carbon dioxide and water vapor in air at room temperature and atmospheric pressure (*90*).

#### 2.2 Guinea pig geometry and thermal boundary conditions

We modeled the guinea pigs as ellipsoids of dimensions consistent with the typical size of adult guinea pigs: semi-axes of 7 cm, 5 cm, and 5 cm. They were lowered by ≈ 1 cm to ensure a lower part contact with cage floor surface for heat exchange, resulting in ≈ 9 cm of vertical height and 14 cm in body length, i.e., body volume of ≈ 733 $cm^3$ and corresponding body mass of ≈ 700-750 g assuming water density. We imposed boundary conditions consisting of a constant heat flux across the guinea pig's body surface, thermally insulating side and top walls (adiabatic boundaries), and a cage floor maintained at constant ambient temperature of $20°C$. A constant heat flux boundary condition was used rather than a fixed surface temperature to better reflect physiologic heat generation (*89*).

#### 2.3 Metabolic heat generation and temperature field

Estimates of guinea pig metabolic heat generation were based on classical measurements by H.H. Kibler, 1947 (*44*) that report resting metabolic rates ranging from ≈ 670 to 820 kcal $m^{-2}$ $day^{-1}$, depending on age and physiological condition: resting metabolism of ≈750 kcal $m^{-2}$ $day^{-1}$ at 2 weeks of age, with peak near ≈ 820 kcal $m^{-2}$ $day^{-1}$ at 10 weeks, and a decline to ≈720 kcal $m^{-2}$ $day^{-1}$ by 7 months. Using the empirical surface-area-to-weight (CGS) correlation (*44*), $SA = 9.85W^{0.64}$, the total metabolic power estimate was between ≈ 2.2 W and 3.1 W for guinea pigs weighing ≈ 0.7-1.2 kg. For the present model, we take 700g of body mass with ellipsoidal surface area of 0.131 $m^2$ and imposed a uniform surface heat flux of 17 W $m^{-2}$, corresponding to a continuous metabolic power of ≈ 2.2 W (Table S3). The resulting temperature field is shown in Supplemental Fig. 8E with maximum temperature variations of $\Delta T \sim 10\,°\mathrm{C}$. Using a characteristic velocity scale $U \sim \sqrt{g\,\beta\,\Delta T\,L}$, and a characteristic length $L \sim 10\ cm$, corresponding to the guinea pig's body size, combined with ambient air properties, yields $U \sim 10$cm/s, which is comparable to flow speeds observed in the simulations (see Supplemental Fig. 3 and 4).

#### 2.4 Guinea pig respiratory physiology and numerical breathing modeling

##### 2.4.1 Physiologic parameters for guinea pig respiration

Physiologic parameters describing guinea pig respiration were drawn from published measurements in adult animals, including respiratory rate, tidal volume, and minute ventilation (*43, 91*).Reported values span a range of animal weights and ages, with adult guinea pigs (≈700-1200 g) exhibiting respiratory rates of approximately 70-104 breaths per minute and tidal volumes ranging from ≈1-4 mL per breath (*45*).

Published measurements of respiratory parameters used to inform the simulations are summarized in Table S3. Reported lung volumes range from approximately 13-39 mL for animals weighing 400 g to 1.3 kg (*46*). These values were used to define representative breathing parameters for an adult guinea pig consistent with the animals used in the transmission experiments.

| | **Admur and Mead (1958)** | **Guyton (1947)** |
|---|---|---|
| | **Mean ± SD** | **Mean (Range)** |
| **Weight** | 220 ± 32 g | 466 g (274-941) |
| **Respiratory rate** | 84 ± 14 breaths/min | 90 (70 - 104) breaths/min |
| **Tidal volume** | 1.68 ± 0.4 mL | 1.75 (1.0 - 4) mL |

| | | |
|---|---|---|
| **Minute ventilation** | 140 ± 30 mL/min | 156 (100 - 382) mL/min |
| **Average exhalation flow rate $Q_{exh}$ based on above studies** | 2.33 ± 0.5 mL/sec | 1.67 - 6.37 mL/sec |
| **Chosen value of exhalation flow rate $Q_{exh}$ in numerics** | $Q_{exh} = 3.75$ mL/sec | |

**Table S3. Physiological parameters of guinea pigs** The values were obtained on 200 normal animals by Admur and Mead (*43*) and 65 animals by Guyton (*91*).

#### 2.4.2 Numerical representation of tidal breathing

To represent tidal breathing in a computationally tractable manner, respiratory exhalation from the donor guinea pig was modeled as a continuous volumetric outflow rather than as discrete breathing cycles. This approximation preserves the correct time-averaged volumetric flux while avoiding prohibitively small time-steps that would be required to resolve individual inhalation-exhalation cycles.

Based on the physiologic ranges summarized above, a representative exhalation flow rate of $Q_{exh}$ = 3.75 mL $s^{-1}$ was selected for the simulations. This corresponds to an exhaled volume of approximately 2.5 mL per breath at a breathing frequency of 90 breaths per minute, which falls within the ranges reported for adult guinea pigs (*43, 91*).

Tidal breathing was implemented as a steady inflow boundary condition at the donor guinea pig mouth, assumed to be circular with a diameter of 1 cm and a fully developed velocity profile. This continuous-flow representation captures the mean respiratory contribution to airflow and tracer transport within the housing system.

#### 2.4.3 Numerical representation of cough emissions

To model episodic cough events, a transient puff-like emission was superimposed on the steady breathing flow. Each cough was represented as a one-second pulse injection of 30 mL of tracer-laden air emitted from the donor guinea pig mouth.

This pulse-based representation captures the short-duration, high-volume nature of cough emissions while remaining computationally feasible. In the simulations, cough events were introduced on top of the steady breathing background flow to isolate the effects of episodic exhalations on airflow structure and tracer transport. Representative cough-puff simulations are provided in Supplemental Movies 6 and 7.

**2.4.4 Flow-regime analysis: relative importance of momentum and buoyancy**

To evaluate the relative importance of momentum and buoyancy in guinea pig respiratory flows, we examined characteristic nondimensional parameters governing exhalation-driven transport.

We first consider the Reynolds number, Re, of the exhaled flow, which quantifies the relative importance of momentum to viscous effects. For a guinea pig tidal exhalation,

$$\mathrm{Re} = \frac{UL}{\nu} \approx \frac{0.048 \times 0.01}{1.5 \times 10^{-5}} \approx 32,$$

where L corresponds to the characteristic mouth opening diameter (d = 1 cm), $U$ is the characteristic exhalation velocity estimated from the volumetric flow rate $U = Q_{\mathrm{exhale}}/A_{\mathrm{mouth}} \sim 0.048$ m/s, and $\nu = 1.5 \times 10^{-5}\ \mathrm{m^2\ s^{-1}}$ is the kinematic air viscosity. Thermal buoyancy induced by guinea pig metabolic heat is characterized by the Grashoff number,

$$\mathrm{Gr} = \frac{g\,\beta\,\Delta T\,L^3}{\nu^2} \approx \mathcal{O}(10^3),$$

using parameters listed in Table S1 and S2 and a characteristic temperature difference of $\Delta T \approx 10^{\circ}$C (Supplemental Fig. 8E).

The relative importance of buoyancy and momentum is captured by the Richardson number,

$$\mathrm{Ri} = \frac{\mathrm{Gr}}{\mathrm{Re}^2} \sim \mathcal{O}(1).$$

Thus, buoyancy and momentum contributions are of comparable magnitude under these conditions, indicating that buoyancy effects must be incorporated to capture the dynamics governing transport of exhaled contaminants and pathogens in the guinea pig transmission systems evaluated here.

### 2.5 Exposure metric and inhalation layer

#### 2.5.1 Simulation procedure for breathing and coughing

To reproduce animal breathing and coughing behaviors, a two-step simulation procedure was used. First, a steady-state simulation with a constant exhalation velocity from the mouth was performed to obtain the baseline airflow and temperature fields associated with breathing. Next, an impulsive inlet flow was applied to represent a short-duration (1 s) cough, and a fully transient simulation was conducted to capture the temporal evolution of airflow, heat, and tracer concentration fields.

#### 2.5.2 Definition of exposure metric

To quantify exposure of the recipient naïve guinea pig to pathogen-like tracers emitted from the donor, we defined a time-averaged tracer concentration, $\bar{c}(t)$, within the inhalation region, as discussed in the main text and illustrated in Fig. 6A.

#### 2.5.3 Definition of the inhalation layer

The inhalation layer was defined as a region of thickness $\delta \approx 1\ cm$ surrounding the recipient guinea pig. The thickness $\delta$ represents a physical estimate of the spatial extent over which the inhalation suction sink significantly influences surrounding airflow, given an inhalation flow rate of $Q_{\mathrm{inh}} \approx 3.75\ \mathrm{mL}/s$ (Table S3).

Air was drawn into the mouth sink from a hemispherical ambient control surface in front of the mouth. Conservation of volume flux over this hemisphere yields a velocity magnitude varying with distance as

$$u(r) = \frac{Q_{\mathrm{inh}}}{2\pi r^2},$$

where, $r$ is the radial distance from the mouth sink (*92*).

**2.5.4 Estimation of inhalation-layer thickness**

An inhalation-layer of thickness $\delta_\alpha$ can be defined as the distance from the guinea pig's mouth at which the local velocity falls below a chosen fraction, $\alpha$, of the driving mouth suction velocity, $U_{\mathrm{mouth}}$, such that

$$u(\delta_\alpha) = \alpha\, U_{\mathrm{mouth}},\ 0 < \alpha < 1.$$

Substituting for $u(r)$ gives

$$\alpha\, U_{\mathrm{mouth}} = \frac{Q_{\mathrm{inh}}}{2\pi\delta_\alpha^2},$$

and therefore

$$\delta_\alpha = \sqrt{\frac{Q_{\mathrm{inh}}}{2\pi\,\alpha\, U_{\mathrm{mouth}}}}.$$

Recalling that $U_{\mathrm{mouth}} = Q_{\mathrm{inh}}/A_{\mathrm{mouth}}$, with inhaling mouth area $A_{\mathrm{mouth}} = \pi(d/2)^2$, this expression reduces to

$$\delta_\alpha = \sqrt{\frac{A_{\mathrm{mouth}}}{2\pi\alpha}} = \sqrt{\frac{\pi(d/2)^2}{2\pi\alpha}} = \frac{d}{2\sqrt{2\alpha}}.$$

For a mouth diameter of $d \approx 1\,\mathrm{cm}$, characteristic inhalation-layer thicknesses corresponding to different velocity thresholds were obtained, for example:

$\delta_{10\%} \approx 1.12\,\mathrm{cm}$, $\delta_{5\%} \approx 1.58\,\mathrm{cm}$, or $\delta_{1\%} \approx 3.54\,\mathrm{cm}$.

##### 2.5.5 Choice of inhalation-layer thickness used in simulations

In the exposure results shown in Fig. 6 and discussed in the main text, we chose a conservative inhalation-layer thickness of $\delta \approx 1.2\,\mathrm{cm}$, corresponding to an approximate 8% decrease in mouth velocity. This choice underestimated absolute exposure while remaining sufficient to capture relative differences in exposure and transmission efficacy across the housing systems considered.

### 3. Negative pressure implementation

#### 3.1 Rationale and assumptions for modeling negative-pressure containment

Guinea pig transmission experiments were conducted under BSL-3 laboratory conditions, in which negative pressure relative to surrounding spaces is used to ensure containment. As shown by the tracer particle experiments in Fig. 3 (main text), not all housing configurations were fully sealed, raising the possibility that facility-level ventilation could alter airflow patterns within the cages. To capture these effects, we explicitly modeled the interaction between ambient negative-pressure ventilation and cage-level leakage pathways.

Given a BSL-3 laboratory ventilation air-change rate (ACH), the volumetric flow rate through the laboratory is

$$Q_{\mathrm{room}} = \frac{\mathrm{ACH} \cdot V_{\mathrm{room}}}{3600},$$

where $V_{\mathrm{room}}$ is the laboratory volume. The corresponding characteristic background airflow velocity was estimated as

$$U \approx \frac{Q_{\mathrm{room}}}{A_{\mathrm{room}}},$$

where $A_{\mathrm{room}}$ is the laboratory cross-sectional area.

We assumed that this background airflow imposed an additional pressure-driven inflow through cage leakage pathways. Based on the geometry and orientation of the housing systems, the front face of the cage was most directly exposed to room-level airflow. Accordingly, we modeled the additional inflow induced by BSL-3 laboratory ventilation as entering primarily through the front leakage strip. All remaining leakage pathways were treated identically to those described under ambient conditions in Section 1.

Because system 2B was mechanically sealed along the front face, no additional inflow due to laboratory background airflow was applied for that configuration. Reduced side leakage strips were retained and modeled as resistive boundaries, as described in Section 1, for consistency across systems.

The dynamic pressure associated with laboratory-scale airflow grazing the exterior of the cage was modeled as a pressure difference across the leakage strips,

$$\Delta p = C_p(1/2)\rho U^2$$

where $C_p$ is an effective surface pressure coefficient relating the local pressure difference at the leakage location to the dynamic pressure of the external flow. Physically, $C_p$ captures the conversion of ambient flow momentum into a static pressure differential across the leakage path and depends on the local orientation of flow impingement, shielding, and degree of flow separation near the opening (*93, 94*).

The resulting air inflow through the leakage strips was modeled using a standard orifice relation,

$$Q_{strip} = C_d A_{strip}\sqrt{2\Delta p/\rho},$$

where $A_{strip}$ is the effective leakage area and $C_d$ is the discharge coefficient of the opening. The discharge coefficient accounts for *vena contracta* formation, flow separation, and viscous losses local to the leakage path, such that the actual inflow is reduced relative to the ideal inviscid orifice prediction (*95-97*). This results in a net inflow of

$$Q_{strip} = C_d A_{strip} U \sqrt{C_p}.$$

Values of $C_p$ and $C_d$ used in the simulations are summarized in Table S4

| **Coefficient** | **Value used in computations** | **Case** |
|---|---|---|
| $C_p$ | 0.3 | GP |
| $C_d$ | 0.6 | Particle, GP |

**Table S4: Values of pressure coefficient and discharge coefficient used to compute inflow from the leakage strip.** For partially normal or oblique impingement typical of grazing indoor flows, representative values are $C_p \approx 0.1 - 0.5$ for short, sharp-edged leakage openings, typical values are $C_d \approx 0.5 - 0.7$ (*93-95*).

| ACH | $U$ (cm/s) | $Q_{\mathrm{strip}}$ (L/min) |
|---|---|---|
| 3 | 0.25 | 0.60 |
| 6 | 0.51 | 1.20 |
| 9 | 0.76 | 1.80 |
| **12** | **1.02** | **2.40** |
| 15 | 1.27 | 3.01 |
| 18 | 1.52 | 3.61 |
| 21 | 1.78 | 4.21 |

**Table S5. Negative-pressure-induced leakage inflow** Estimated inflow through the front leakage strip induced by room-level negative pressure in BSL-3 laboratory.

Laboratory air-change rate (ACH) is given with corresponding characteristic room velocity $U$ (cm $s^{-1}$), inferred from the total ventilation flow rate and the lab cross-sectional area. Resulting leakage-strip inflow rate $Q_{\text{strip}}$ (L $min^{-1}$), computed using a pressure-coefficient-based orifice model with $C_p = 0.3$ and discharge coefficient $C_d = 0.6$. The reported $Q_{\text{strip}}$ for ACH 12 is the inflow rate imposed at the front leakage strip in this study's numerical computations to capture BSL3 negative-pressure conditions.

**Table S5** summarizes estimated leakage inflow rates $Q_{\text{strip}}$ for a range of laboratory ACH values. Estimates are based on the BSL-3 facility housing the guinea pigs (floor area 179 ft²; ceiling height 10 ft). Although BSL-3 laboratories require a minimum ACH of 6, facilities housing animals typically operate at higher values (10-15 ACH) (*98, 99*). For ACH = 12, the estimated leakage inflow is $Q_{\text{strip}} \approx 2.4\ \text{L/min}$ which was used as the imposed front-leakage inflow rate in the transmission-system simulations to represent BSL-3 containment conditions.

**3.2 Particle simulations in negative ambient pressure containment**

In addition to modeling BSL-3 laboratory containment relevant to guinea pig transmission experiments, we separately simulated the negative-pressure chamber used for tracer particle-only experiments, which provided an independent benchmark for leakage-driven airflow under controlled conditions.

For the tracer particle-only transmission measurements (Fig. 3) experiments were conducted in a dedicated negative-pressure chamber rather than in a BSL-3 animal facility. We therefore modelled this chamber separately to reproduce the flow conditions imposed during particle release and sampling experiments and to benchmark leakage-driven airflow under controlled conditions.

In the experiments, the external chamber was maintained at a prescribed static pressure offset of $\Delta p = -0.03 \text{ in } H_2O = -7.5 \text{ Pa}$ relative to the surrounding laboratory using HEPA-filtered air exchange at the chamber boundary. To estimate the characteristic chamber-scale airflow induced by this pressure difference, we used a pressure-based scaling analysis. Treating the chamber-level air exchange as an effective orifice of characteristic diameter $D_{\text{port}} = 15\, cm$, the volumetric flow rate through the chamber was estimated using a Bernoulli-type relation,

$$Q_{outer} \sim C_{d,port} A_{port} \sqrt{\frac{2|\Delta p|}{\rho}},$$

where air density is $\rho = 1.2 \text{ kg m}^{-3}$, and $C_{d,\text{port}}$ is an effective discharge coefficient (see Table S4).

Using this scaling estimated a chamber-scale volumetric flow rate of $Q_{\text{outer}} \approx 1900 - 2700$ L $\text{min}^{-1} \approx 68\text{-}96 \text{ CFM}$ where CFM indicates flow rate in cubic feet per minute. Given the cross-sectional area of the chamber (24 x 24 $\text{in}^2$), this corresponds to a characteristic grazing airflow velocity of approximately $U \approx 8\text{-}12 \text{ cm s}^{-1}$ along the cage exterior.

To translate this chamber-scale background flow into an effective inflow through cage leakage pathways, we applied the same leakage-strip modeling framework described in Section 3.1. Substituting the estimated velocity range into the leakage inflow relation and assuming a surface pressure coefficient in the range $C_p \approx 0.05 - 0.30$ yields an expected inflow through the front leakage strip of $Q_{\text{strip}} \approx 8\text{-}29 \text{ L min}^{-1}$.

For the tracer particle-only simulations reported in the manuscript, we selected a representative value within this range, $Q_{\text{strip}} \approx 12 \text{ L min}^{-1}$ as a conservative middleground estimate. This value was imposed as the inflow boundary condition at the front leakage strip in the numerical simulations used to reproduce the particle-only experimental conditions shown in Fig. 3.

### Supplemental Figure 1

A

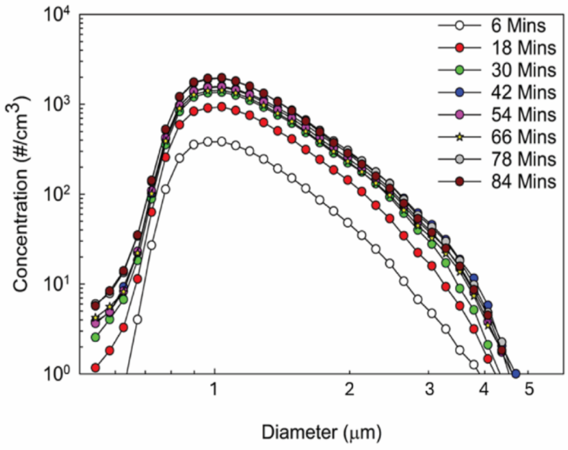

B

| Chemical Compound | Concentration (g/L) |
|---|---|
| $CaCl_2$ (anhydrous) | 0.2 |
| KCl | 0.4 |
| $MgSO_4$ (anhydrous) | 0.0977 |
| NaCl | 6.8 |
| $NaH_2PO_4.H_2O$ | 0.14 |
| $NaHCO_3$ | 2.2 |
| L-Arginine.HCl | 0.1264 |
| L-Cystine .2HCl | 0.0312 |
| L-Glutamine | 0.292 |
| L-Histidine.HCl.$H_2O$ | 0.0419 |
| L-Isoleucine | 0.0525 |
| L-Leucine | 0.0525 |
| L-Lysine. HCl | 0.0725 |
| L-Methionine | 0.015 |
| L-Phenylalanine | 0.0325 |
| L-Threonine | 0.0476 |
| L-Tryptophan | 0.01 |
| L-Tyrosine.2Na.2$H_2O$ | 0.0519 |
| L-Valine | 0.0468 |
| D-Calcium pantothenate | 0.001 |
| Choline chloride | 0.001 |
| Folic acid | 0.001 |
| i-Inositol | 0.002 |
| Nicotinamide | 0.001 |
| Pyridoxine. HCl | 0.001 |
| Riboflavin | 0.0001 |
| Thiamine.HCl | 0.001 |
| D-Glucose | 1 |
| Phenol red.Na | 0.01 |

**Supplemental Figure 1: Particle generation within cage systems.** (A) Concentration equilibrium within cage systems from 6 to 84 minutes of continuous generation. (B) chemical composition of EMEM used for particle generation.

Supplemental Figure 2

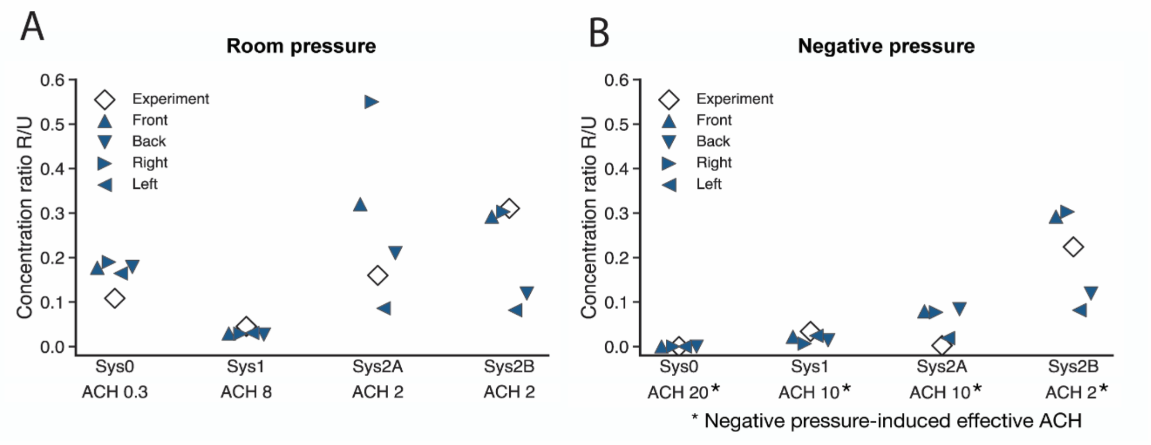

**Supplemental Figure 2: Sensitivity of simulated concentration ratios to probe location.** Computed recipient-to-upstream tracer concentration ratios, R/U, evaluated at individual sampling probe locations. Results are shown for ambient lab-pressure conditions (left) and negative chamber-pressure conditions (right). Each marker indicates a distinct probe location in the recipient side in the guinea pig BZ, capturing the reasonable spatial variability in placement of a probe for tracer concentration sampling.

Supplemental Figure 3

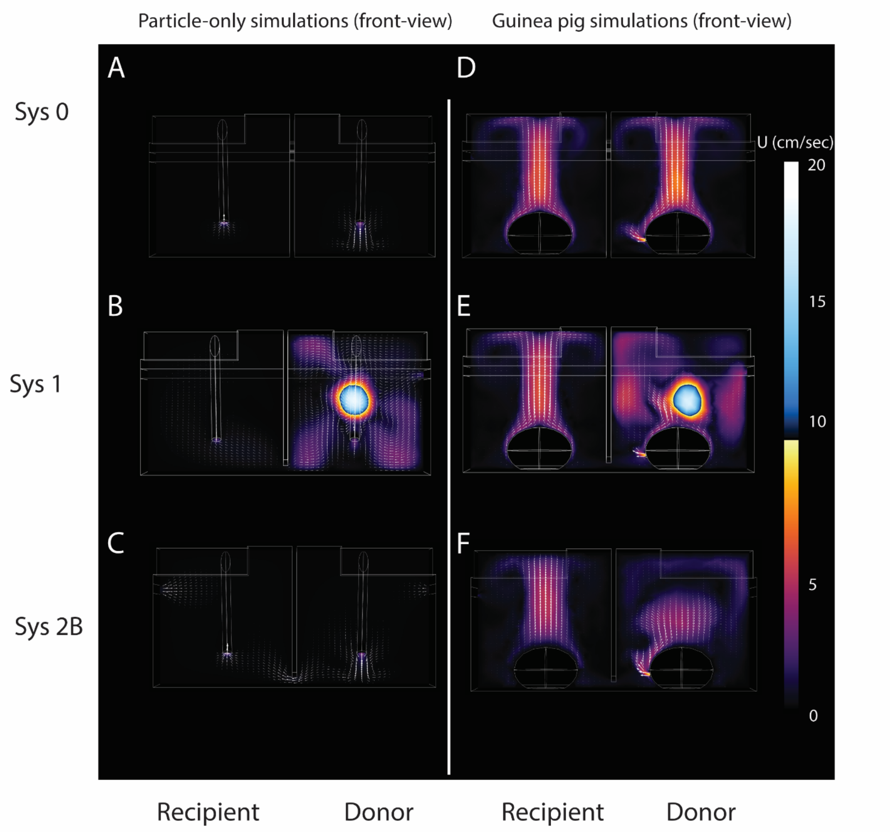

**Supplemental Figure 3: Effect of buoyancy on airflow structure across housing systems.** Vertical midplane concentration slices comparing particle-only simulations (A,B, and C) and guinea pig-resolved simulations with body heat-induced buoyancy (D, E, and F) for systems 0, 1, and 2B. Background color indicates the magnitude of the three-dimensional velocity field, while in-plane velocity vectors (u,v) are overlaid to reflect flow field within the slice. Particle-only snapshots (left) correspond to steady-state solutions obtained in the absence of animals. Guinea pig-resolved snapshots (right) correspond to steady-breathing regime, prior to cough event, and include buoyancy effects arising from animal body heat.

Supplemental Figure 4

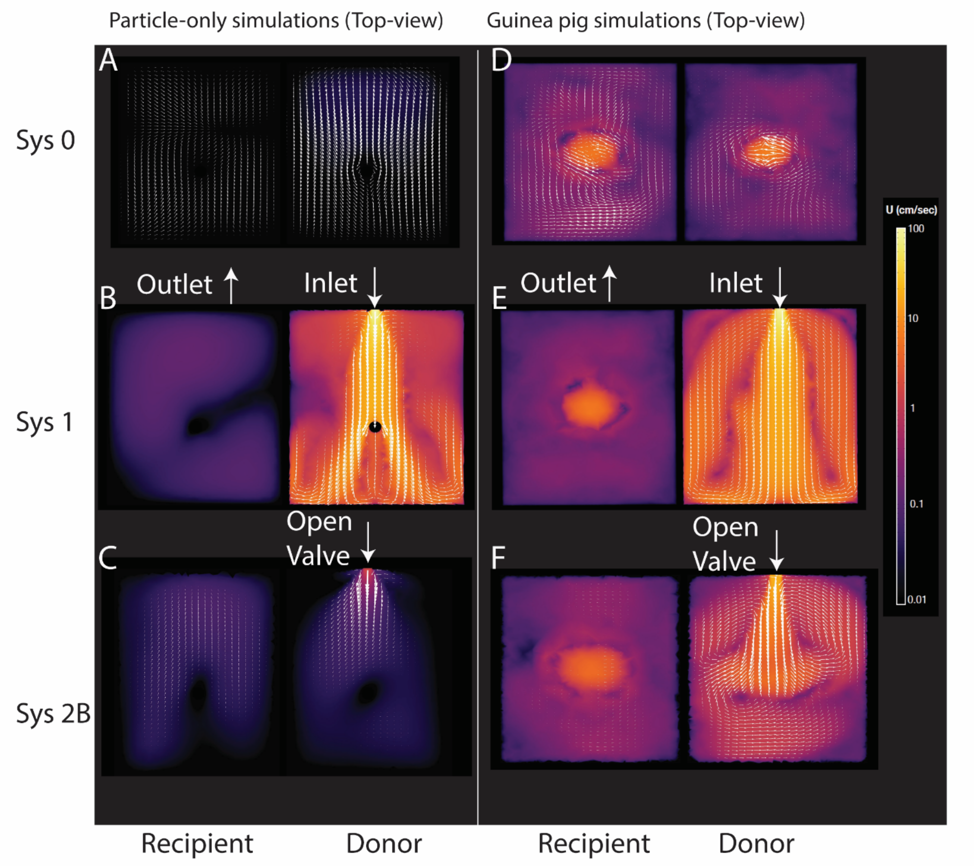

**Supplemental Figure 4: Effect of buoyancy on airflow structure across housing systems.** Horizontal midplane slices comparing particle-only simulations (A, B, and C) and guinea pig simulations including body heat-induced buoyancy (D,E, and F) for systems 0, 1, and 2B. Background color indicates the magnitude of the three-dimensional velocity field, while in-plane velocity vectors (u,w) are overlaid to illustrate flow field within the slice. Other details are same as supplemental Fig. 3. A circular patch of higher velocity magnitude mainly present due to flow velocity normal to the plane is due to buoyancy. Horizontal slices are taken to be in the middle of the cage, around $\approx 7cm$, corresponding to the upper part of guinea pig BZ.

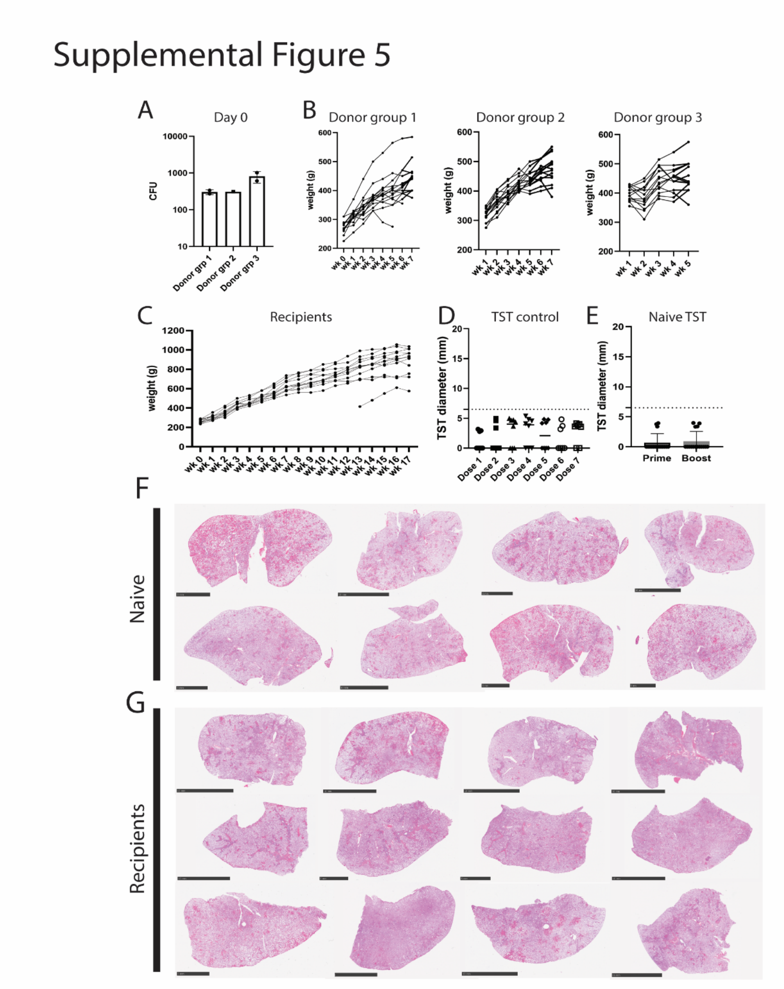

**Supplemental Figure 5: Animal characteristics of transmission study**. (A) Lung CFU directly after aerosol infection (day 0) for donor groups 1,2, and 3. (B) Weight tracking of donor guinea pigs over 7 weeks (group 1 and 2) and 5 weeks for group 3. (C) Weight tracking of recipient guinea pigs throughout 17-week exposure period. (D) TST measurement in TST control guinea pigs given 7 doses of tuberculin. (E) TST measurement in naïve guinea pigs given first and second dose of tuberculin. Dashed line denotes cut-off (6.5 mm) for positive responses. (F) Hematoxylin and eosin staining of lung sections from 8 naïve animals used for histopathology analysis. (G) Lung sections from 12 recipient animals stained with hematoxylin and eosin for histology analysis.

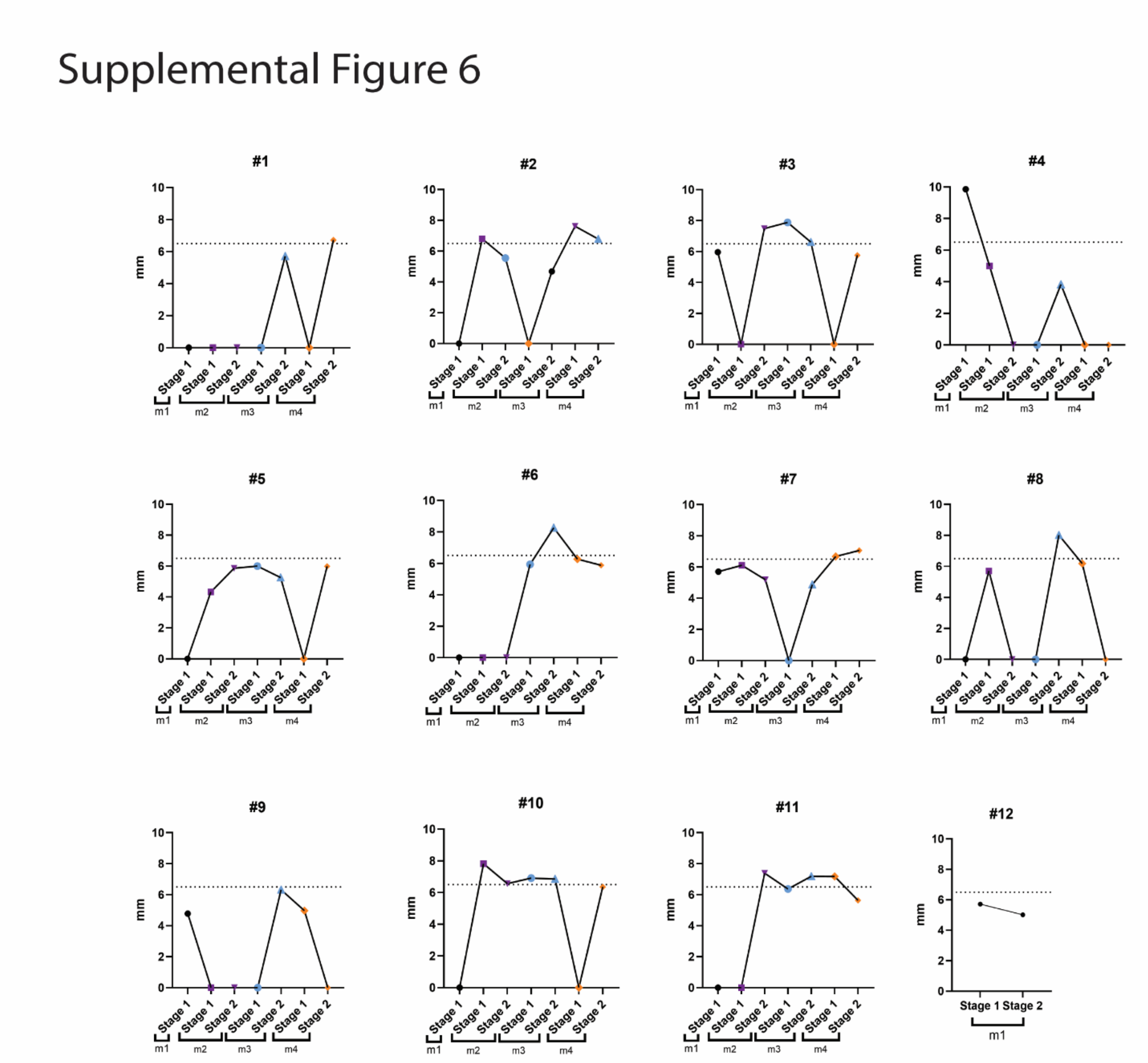

**Supplemental Figure 6: TST measurements of individual recipient guinea pigs.** TST measurements for each recipient guinea pig (#1-12) across the exposure period. Any values above 6.5 mm (dashed line) are considered positive. Recordings include the first (stage 1) and second (stage 2) dose of tuberculin.

Supplemental Figure 7

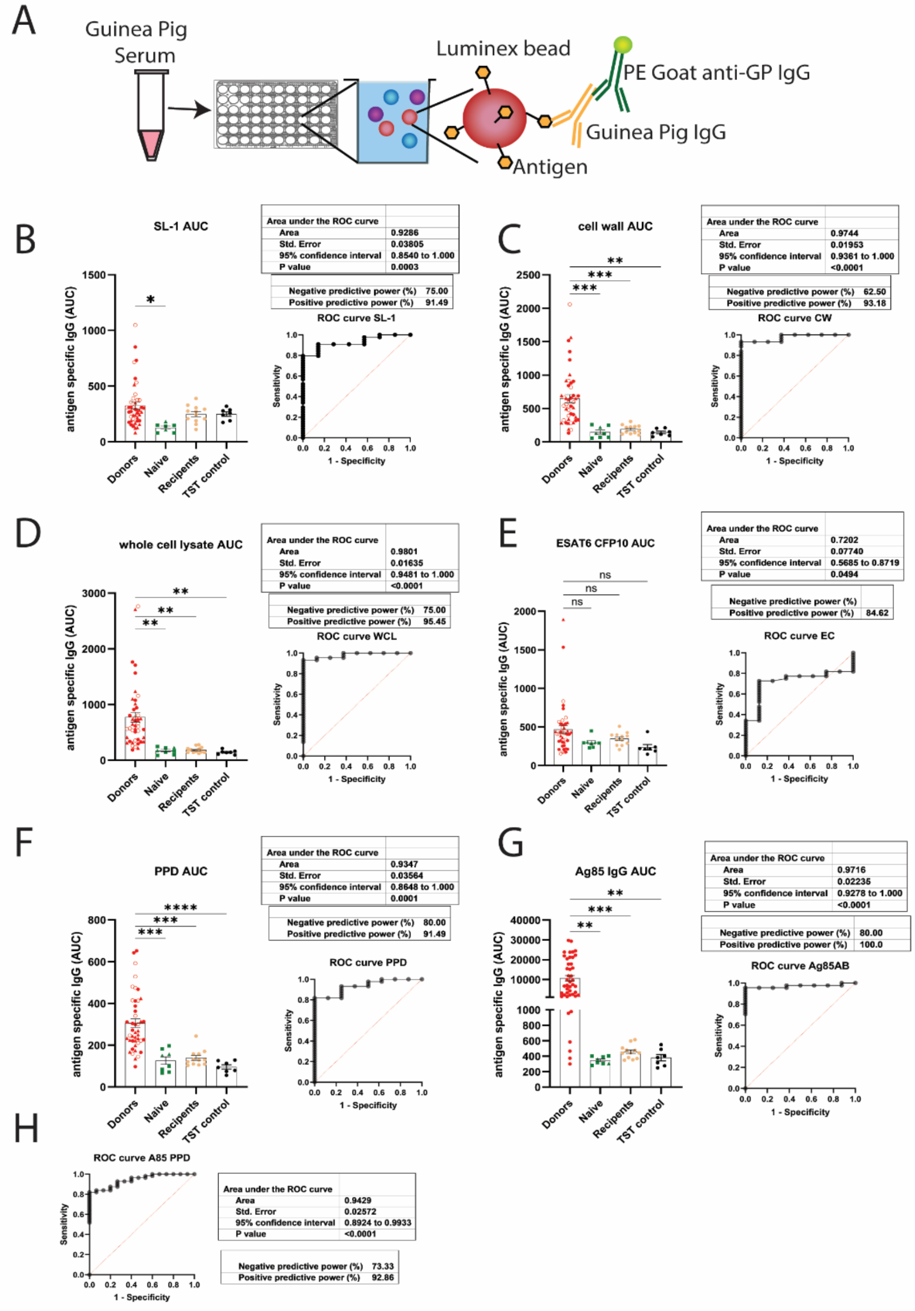

**Supplemental Figure 7: Serum antibody levels of guinea pigs.** (A) experimental design of bead-based serum antibody quantification. (B) Area under the curve (AUC) of SL-1 specific IgG for donor, naïve, recipient, and TST control groups. Receiver operation characteristic (ROC) curve predictive power (%) for both positive and negative values based on SL-1 antigen specific IgG. (C-G) AUC and ROC curves for cell wall, whole cell lysate, ESAT6 CFP10, PPD, and Antigen 85B specific IgG. (H) ROC curve for combined antigen 85B and PPD IgG.

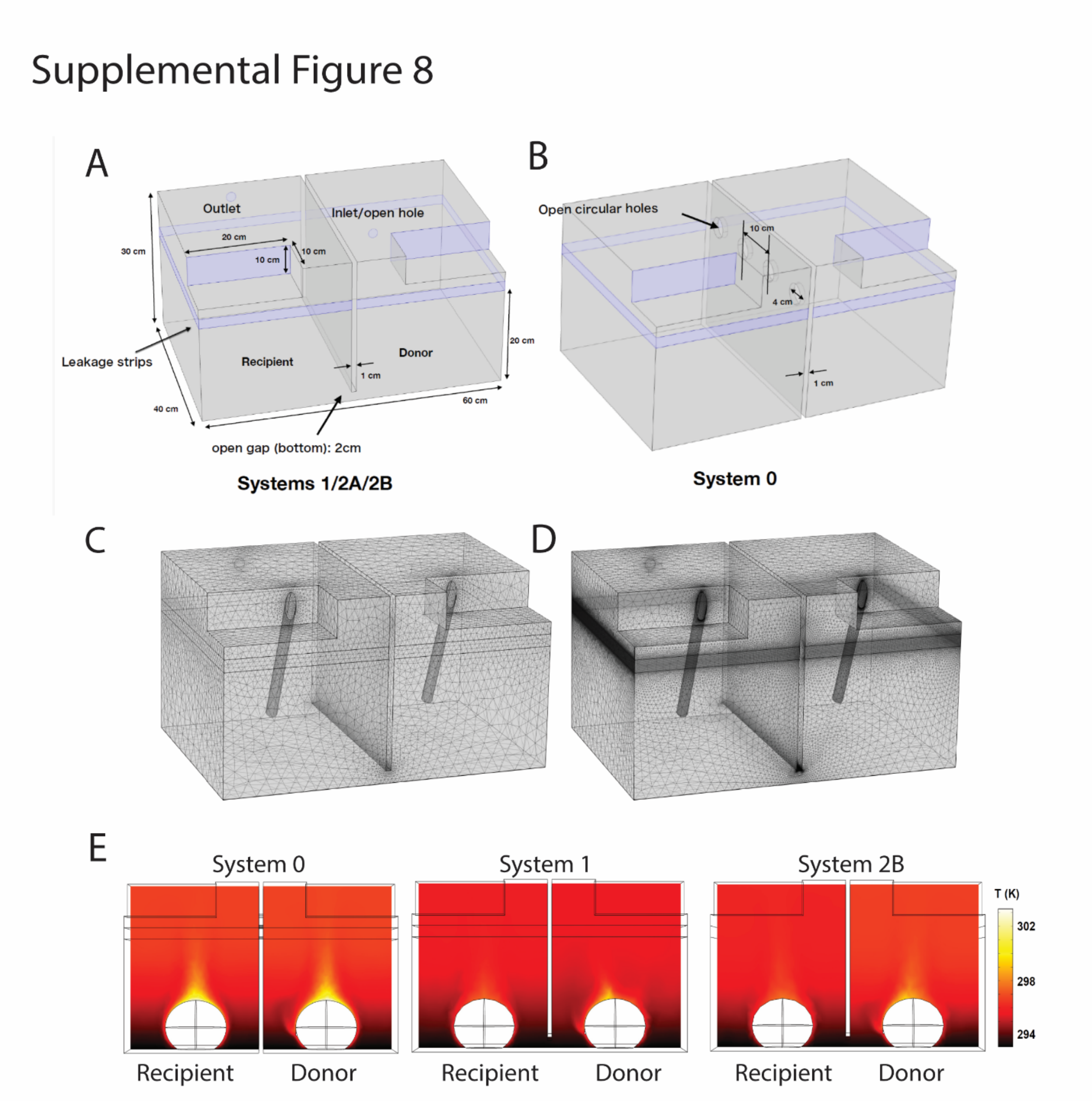

**Supplemental Figure 8:** (A, B) Domain dimensions used for computations recapitulating the experimental system**.** Left: System 1/2A/2B and Right: System 0. The diameter of inlet/open and outlet holes are 2 cm. The width of the separator is 1 cm and the open gap at the bottom is 2 cm. Leakage strips are 2 cm wide for all systems. To closely mimic experiments, for system 2B, the front and back strips are sealed and side strips are given a flow resistance modeling thin leakage strip. For system 1, the valve on the Donor side acts as an inlet of volumetric flow rate 8.8 L/min while it is left as an open boundary for systems 2A and 2B. (C,D) Two mesh sizes for system 2A particle-only simulations. The left mesh shows a coarse mesh with mesh refinement factor of h=1 as defined in the **Table S2** . The right mesh is a finer mesh for h=0.125 and the one used in the simulations. Both meshes are generated using identical growth-rate, curvature, and corner-refinement settings, with refinement applied consistently in the bulk domain, leakage strips, inlet-outlet regions, and probe region as described in **Table S2** (E) Temperature fields. The temperature variation within the domain is of order 10° C, with the highest temperatures occurring near the upper surface of the guinea pig body, reaching approximately 300 K ≈ 27° C to maintain physiological heat flux. The guinea pig surface temperature in systems 1 and 2B is lower than system 0, due to additional convective cooling associated with ventilation airflow in these cage systems.

**Supplemental Movie 1**: **Airflow smoke test.** Unidirectional airflow of system 1,2A and 2B cages as determined by smoke injection into the donor compartment and passage into the recipient compartment.

**Supplemental Movie 2: Cross-sectional tracer concentration across cage configurations.** Tracer concentration fields on successive cross-sectional slices spanning the cage width for all housing systems under ambient lab pressure conditions, providing a dynamic extension of the mid-plane slices shown in Fig. 5 E-F. The slice plane translates across the domain volume.

**Supplemental Movie 3: Cross-sectional tracer concentration across cage configurations under negative pressure.** Tracer concentration fields on successive cross-sectional slices spanning the cage width for all housing systems under negative pressure conditions.

**Supplemental Movie 4: Visualization of airflow topology using frozen streamlines.** Airflow pathways originating from the donor side using frozen streamlines computed from the simulated velocity field. Streamlines are colored by normalized tracer concentration and are held fixed at a selected time instant and therefore represent the developed flow pattern; the movies do not depict transient flow evolution. Passive tracer particles are periodically injected at the streamline starting locations and animated along these frozen streamlines solely to visualize flow topology and transport pathways. Particle motion is shown for a total physical duration of 1000 s, with particles injected every 100 s. For visualization purposes, this duration is mapped to a 50 s movie, such that movie time scales linearly with physical time and can be directly compared across systems (e.g., 25 s in one movie corresponds to the same physical time as 25 s in another). Right panel shows the guinea pig cage system, with streamlines seeded at the donor guinea pig mouth. Streamlines are frozen at an instant 0.1 s after the onset of a cough event (the cough duration is 1 s), capturing the instantaneous flow field generated by respiratory momentum and buoyancy. Left panels show the corresponding particle-only simulation, with streamlines seeded at the donor inlet tube under steady-state conditions.

**Supplemental Movie 5: Visualization of airflow topology in frozen streamlines:** Illustration of the airflow pathways originating from the donor side using frozen streamlines for the guinea pig cage system, with streamlines seeded at the donor guinea pig mouth. Streamlines are frozen at 0.1 s post-cough onset (1s cough duration), capturing the instantaneous flow field generated by respiratory momentum and buoyancy. The streamlines are colored by normalized tracer concentration. The left panel shows the streamline animation in full system, while the right panel shows the zoomed-in recipient guinea pig, i.e. reflecting its exposure.

**Supplemental Movie 6:** Movie showing mid-plane slice coloured by the three-dimensional velocity magnitude, with vectors indicating the in-plane velocity components. A cough event is initiated at t = 0 and lasts for 1 s, following which steady breathing starts again. During the 1 sec of cough event a total volume of 30 mL is expelled from the donor guinea pig mouth. The cough is injected on an already developed baseline flow field generated by steady breathing. The movie shows the subsequent evolution of the flow field for 5 s following the onset of cough.

**Supplemental Movie 7:** Mid-plane slice with colour-scale indicating concentration field, normalized by the donor emission concentration. Contaminated air with tracer concentration injected from the donor guinea pig mouth. A cough event is initiated at t = 0 and lasts for 1 s, during which a total volume of 30 mL is expelled. The cough is imposed on an already developed baseline tracer field generated by steady breathing, after which steady breathing resumes. The movie shows the subsequent evolution of the tracer concentration field for 5 s following the onset of the cough.
